# Discovery and engineering of the antibody response against a prominent skin commensal

**DOI:** 10.1101/2024.01.23.576900

**Authors:** Djenet Bousbaine, Katherine D. Bauman, Y. Erin Chen, Victor K. Yu, Pranav V. Lalgudi, Arash Naziripour, Alessandra Veinbachs, Jennie L. Phung, Tam T.D. Nguyen, Joyce M. Swenson, Yue E. Lee, Alex Dimas, Sunit Jain, Xiandong Meng, Thi Phuong Thao Pham, Aishan Zhao, Layla Barkal, Inta Gribonika, Koen K.A. Van Rompay, Yasmine Belkaid, Christopher O. Barnes, Michael A. Fischbach

## Abstract

The ubiquitous skin colonist *Staphylococcus epidermidis* elicits a CD8^+^ T cell response pre-emptively, in the absence of an infection^1^. However, the scope and purpose of this anti-commensal immune program are not well defined, limiting our ability to harness it therapeutically. Here, we show that this colonist also induces a potent, durable, and specific antibody response that is conserved in humans and non-human primates. A series of *S. epidermidis* cell-wall mutants revealed that the cell surface protein Aap is a predominant target. By colonizing mice with a strain of *S. epidermidis* in which the parallel β-helix domain of Aap is replaced by tetanus toxin fragment C, we elicit a potent neutralizing antibody response that protects mice against a lethal challenge. A similar strain of *S. epidermidis* expressing an Aap-SpyCatcher chimera can be conjugated with recombinant immunogens; the resulting labeled commensal elicits high titers of antibody under conditions of physiologic colonization, including a robust IgA response in the nasal mucosa. Thus, immunity to a common skin colonist involves a coordinated T and B cell response, the latter of which can be redirected against pathogens as a novel form of topical vaccination.

## INTRODUCTION

The adaptive immune response to a pathogen has a clear goal: limiting replication and dissemination to prevent the host from dying. By comparison, the adaptive immune response to microbial colonists is puzzling. Three of its features are notable: its tone is determined by the microbial elicitor^1–4^; it is antigen-specific^5–11^, just like an anti-pathogen response; and it is pre-emptive—it occurs in the absence of infection or inflammation, across an intact barrier. Given that microbial residents of the gut, skin, and nasal cavity are thought to be generally mutualistic, what is the purpose of this response?

We chose to pursue this question in the context of the ubiquitous skin colonist *Staphylococcus epidermidis*. Certain strains of *S. epidermidis* induce antigen-specific CD8^+^ T cells in mice and non-human primates^1,12^. We recently showed that by colonizing mice with strains of *S. epidermidis* engineered to produce tumor antigens, we could elicit a potent T cell response against a distant tumor, showing that colonist-specific T cells are not simply homeostatic—they are capable of traveling systemically and sufficiently licensed to kill cells in an immunologically ‘cold’ tumor^13^.

These data suggest that the immune response to *S. epidermidis* is similar to the anti-pathogen response in that it is directed (at least in part) at limiting replication and dissemination, but distinct in that it occurs pre-emptively. We reasoned that these findings are consistent with a program devoted to ‘anticipatory’ immunity, in which the cost of an adaptive immune response to a colonist is pre-paid in order to be prepared for an inevitable barrier breach in the form of a nick, cut, or scrape.

We were intrigued by this model but questioned whether it was complete. Most adaptive immune responses involve the coordinated action of T cells and B cells, but the role of the latter was undefined here. Three pieces of evidence suggested the plausibility of an antibody response against a skin commensal: (*i*) Electron micrographs of human skin swabs have shown that antibodies coat bacterial commensals^14^. (*ii*) Humans colonized with *Staphylococcus aureus* have a higher titer of anti-*S. aureus* antibodies than non-carriers or intermittent carriers^15^, indicating that *S. aureus* may elicit a B cell response during colonization, not just infection. (*iii*) A series of studies have revealed IgA and IgG responses to colonists of the gut^6,16–19;^ these commensal-specific antibodies contribute to barrier function, prevent dissemination^20^, and promote symbiotic relationships with the microbiome^18^. Thus, we reasoned that the host may respond to *S. epidermidis* colonization with antigen-specific B cells, and if so, this would reveal a new component of the immune response to skin colonists.

Here, we show that colonization with *S. epidermidis* elicits a systemic antibody response that is potent, durable, and specific. A predominant target of this response is the large cell-wall-anchored protein Aap, which belongs to a superfamily that is widely distributed among strains of *S. epidermidis*^21^. Mice colonized by a strain of *S. epidermidis* in which the parallel β-helix domain of Aap is replaced by tetanus toxin fragment C are fully protected against a lethal challenge with tetanus toxin. By engineering *S. epidermidis* to express an Aap-SpyCatcher chimera, we can conjugate its surface with recombinant immunogens; the resulting labeled commensal elicits high titers of neutralizing antibody under physiologic conditions of colonization. Thus, pre-emptive immunity against *S. epidermidis* involves the coordinated action of T cells and B cells, and engineered skin colonists are a simple but powerful way to elicit an antibody response against unrelated pathogens.

### *S. epidermidis* induces a potent B cell response

First, we sought to determine whether the immune response to *S. epidermidis* involves B cells in addition to T cells. We chose to study *S. epidermidis* NIHLM087 (LM087), a primary isolate from a healthy donor that elicits antigen-specific CD8^+^ and CD4^+^ T cells under conditions of physiologic colonization^1,12^. We also included *S. epidermidis* NIHLM088 (LM088), a strain isolated from the same donor and body site as LM087 but that does not induce CD8^+^ T cells^12^. We colonized the head of unprepped, conventionally housed C57BL/6 mice gently, using a cotton swab. These conditions do not yield an infection, barrier breach, or inflammation; instead, they simulate physiologic microbial colonization of a host. After colonizing mice for 6 weeks, we harvested serum and measured *S. epidermidis*-specific antibodies by ELISA against immobilized cells of the bacterial colonist. Colonized mice had higher levels of *S. epidermidis-*specific IgG in the serum than uncolonized controls; LM088 induced particularly high titers (**Fig. 1a**). Similar results were obtained by flow cytometry, indicating that the binding of serum antibodies to *S. epidermidis* is not an artifact of the assay (**fig. s1**).

**Figure 1:**
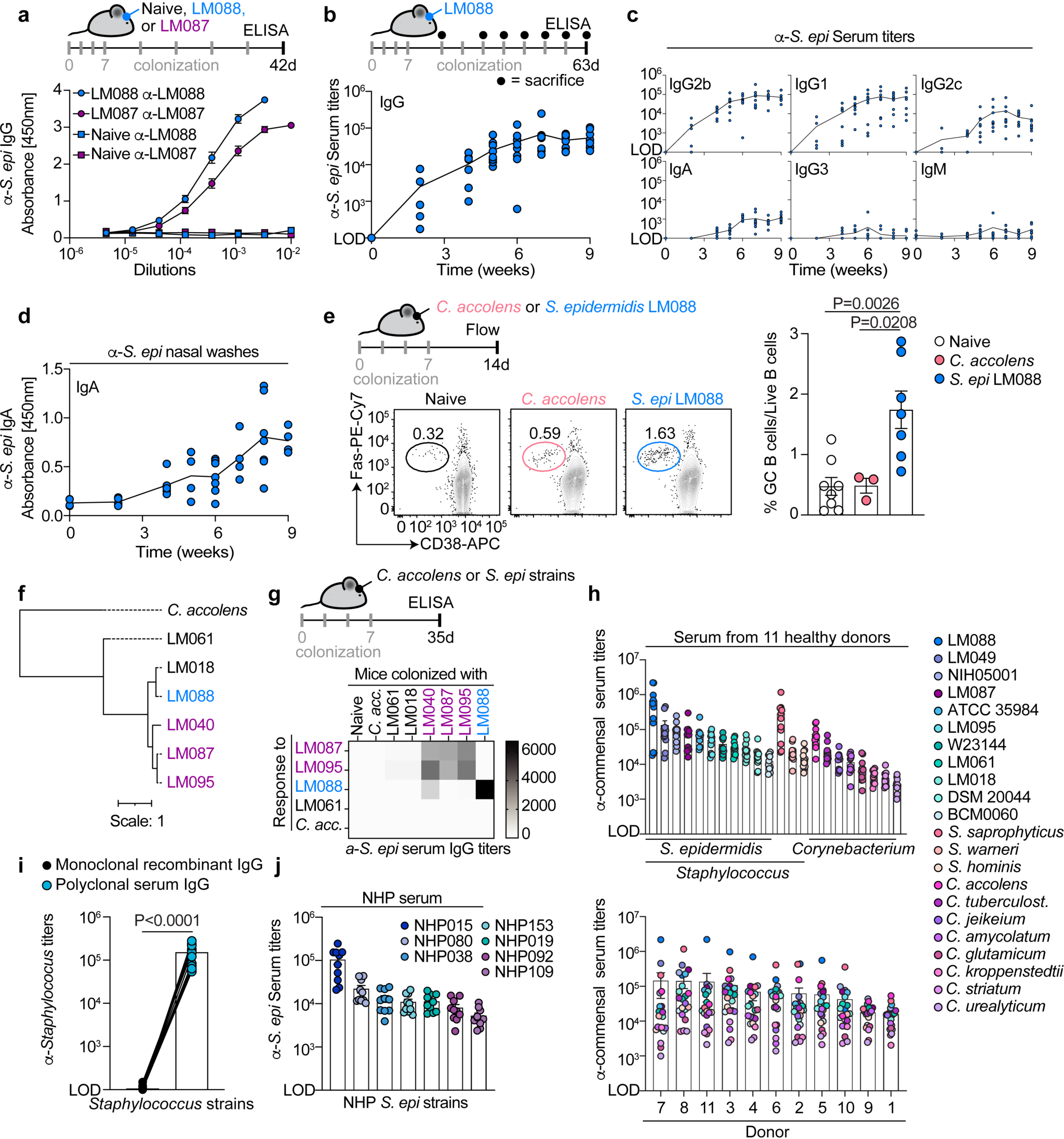
Colonization with *Staphylococcus epidermidis* induces a potent B cell response. (**a**) Specific pathogen-free (SPF) mice were colonized with *S. epidermidis* strain LM087 or LM088. After 6 weeks, serum was harvested and antibody titers measured against the colonist by ELISA (n=23-24/group, three independent experiments pooled). (**b-d**) SPF mice were colonized with LM088 for 2-9 weeks; serum and nasal washes were tested for the presence of α-LM088 antibodies by ELISA. The graphs show all of the mice that were analyzed (n=5-9/timepoint, two independent experiments pooled). (**e**) Frequency of germinal center B cells among live B cells (Live^+^CD19^+^CD4^-^) in the draining cervical lymph nodes (dLNs) of one representative mouse/group (left) and all mice analyzed (right) (n=3-8 mice/group, two independent experiments pooled). (**f**) Phylogenetic analysis of a set of *S. epidermidis* strains. (**g**) SPF mice were colonized with the indicated strains; five weeks later, the serum was tested for α-*S. epidermidis* antibodies and their strain specificity (n=4-8/group, two independent experiments pooled). (**h**) Bacterial IgG ELISA against a panel of *Staphylococcus* and *Corynebacterium* human skin isolates using the serum of 11 healthy human donors. The data are displayed by bacterial strain (top) or by donor (bottom). All donors were tested in one experiment. (**i**) Bacterial IgG ELISA against the panel of *Staphylococcus* isolates used in (**h**) with either an unrelated monoclonal antibody or polyclonal antibodies isolated from human sera. We assumed that the concentration of IgG in human serum was 16 mg/ml^53^ and serially diluted the antibody mixture similarly to (**h**). (**j**) Bacterial IgG ELISA using the serum from 10 non-human primate (NHP) donors against *S. epidermidis* strains isolated from their skin. P-values were calculated using one-way ANOVA (**e**) with Tukey’s multiple comparisons test and paired Student t test (**i**). LOD: limit of detection. All graphs show mean +/− SEM.

Next, we analyzed the magnitude of the antibody response over time. Mice were colonized with LM088 by gentle topical application; at each timepoint a group of mice was euthanized, serum was collected, and antibody titers were measured (**Fig. 1b**). We observed increasing levels of LM088*-*specific IgG that reached ∼2 x 10^4^ by six weeks and remained steady for the duration of the nine-week experiment (**Fig. 1b**). Serum antibodies were dominated by IgG1 and IgG2b, with lower levels of other isotypes (IgG2b>IgG1>IgG2c>IgA) (**Fig. 1c**). Notably, we observed increasing levels of *S. epidermidis*-specific IgA in the nares (**Fig. 1d**), though it is unclear whether this was an indirect consequence of skin colonization or a direct response to incidental colonization of the nares.

Having observed a systemic B cell response involving class-switched antibodies, we reasoned that a germinal center might be present in the draining lymph nodes. Two weeks after the initiation of colonization, we examined B cells in the skin-draining lymph nodes for markers of germinal center induction. We found that LM088 elicited germinal center B cells, whereas *Corynebacterium accolens*—a skin colonist that induces ψ8 T cells^22^—did not (**Fig. 1e**).

Antibody responses can be highly specific. We tested the cross-reactivity of serum from mice colonized by a single strain of *S. epidermidis* against other strains. Serum IgG against *S. epidermidis* was generally strain-specific, though we observed cross-reactivity between members of the same clade (LM087, LM040 and LM095). There was no cross-reactivity with the unrelated skin commensal *C. accolens* (**Fig. 1f-g**). These results are consistent with the view that anti-*S. epidermidis* antibodies recognize a combination of strain-specific and shared antigens that are phylogenetically restricted. Thus, physiologic colonization by *S. epidermidis* yields a potent, durable, and specific antibody response.

### Serum from humans and non-human primates contains antibodies against *S. epidermidis*

Since *S. epidermidis* is a ubiquitous colonist of human skin, we next asked whether the B cell response is conserved in humans. We screened serum samples from eleven healthy human donors against a panel of human *S. epidermidis* isolates (**Fig. 1h**). We included three additional *Staphylococcus* species that are prevalent members of the skin microbiome (*S. saprophyticus, S. warneri, S. hominis*) and a small set of *Corynebacterium* strains, also ubiquitous human commensals^23^. We found very high levels of antibody against every strain of *S. epidermidis,* especially LM088 (titer = 1 x 10^4^ – 2 x 10^6^), and against *S. saprophyticus*, a colonist of the urogenital tract (**Fig. 1h**, top). Antibodies against *Corynebacterium* species were present but at a lower titer.

Every human serum sample had high titers of antibody against *S. epidermidis* (1 x 10^4^ – 2 x 10^5^) (**Fig. 1h**, bottom). To ensure that the binding of serum antibodies to *Staphylococcus* was not due to the presence of an antibody-binding protein on the cell surface (e.g., *S. aureus* protein A), we tested the binding of monoclonal antibodies (specific for unrelated target) against all of our commensal *Staphylococcus* isolates by ELISA. We did not detect any binding, indicating that antibodies in serum do not bind non-specifically to skin isolates of *Staphylococcus* (**Fig. 1i**). In contrast, a commercial preparation of polyclonal IgG from human serum contains antibodies that bind to *Staphylococcus* strains (**Fig. 1i**), supporting the notion that the human repertoire contains antibodies specific for commensal *Staphylococcus*. Similarly, we observed high titers of IgG in the serum of 10 rhesus macaques against *S. epidermidis* strains isolated from the skin of the same animals (**Fig. 1j**). Although more experimentation will be needed to prove that these antibodies are specific for *S. epidermidis* (and not, e.g., polyreactive against multiple bacterial species), these data suggest the possibility that the antibody response against *S. epidermidis* is conserved and highly potent in humans and non-human primates.

### The predominant target of the antibody response is Aap

Next, we sought to identify the molecular target of the antibody response against *S. epidermidis*. Given that it was observed in ELISA and flow cytometry analyses of intact bacterial cells (see **Fig. 1a and fig. s1**), we hypothesized that the target was a molecule on the surface of the bacterium. We started by incubating wild-type LM088 with serum from LM088-colonized mice and imaging the cells by electron microscopy. We observed antibody labeling of the cell wall (**Fig. 2a**), consistent with the view that the target(s) are surface molecules.

**Figure 2:**
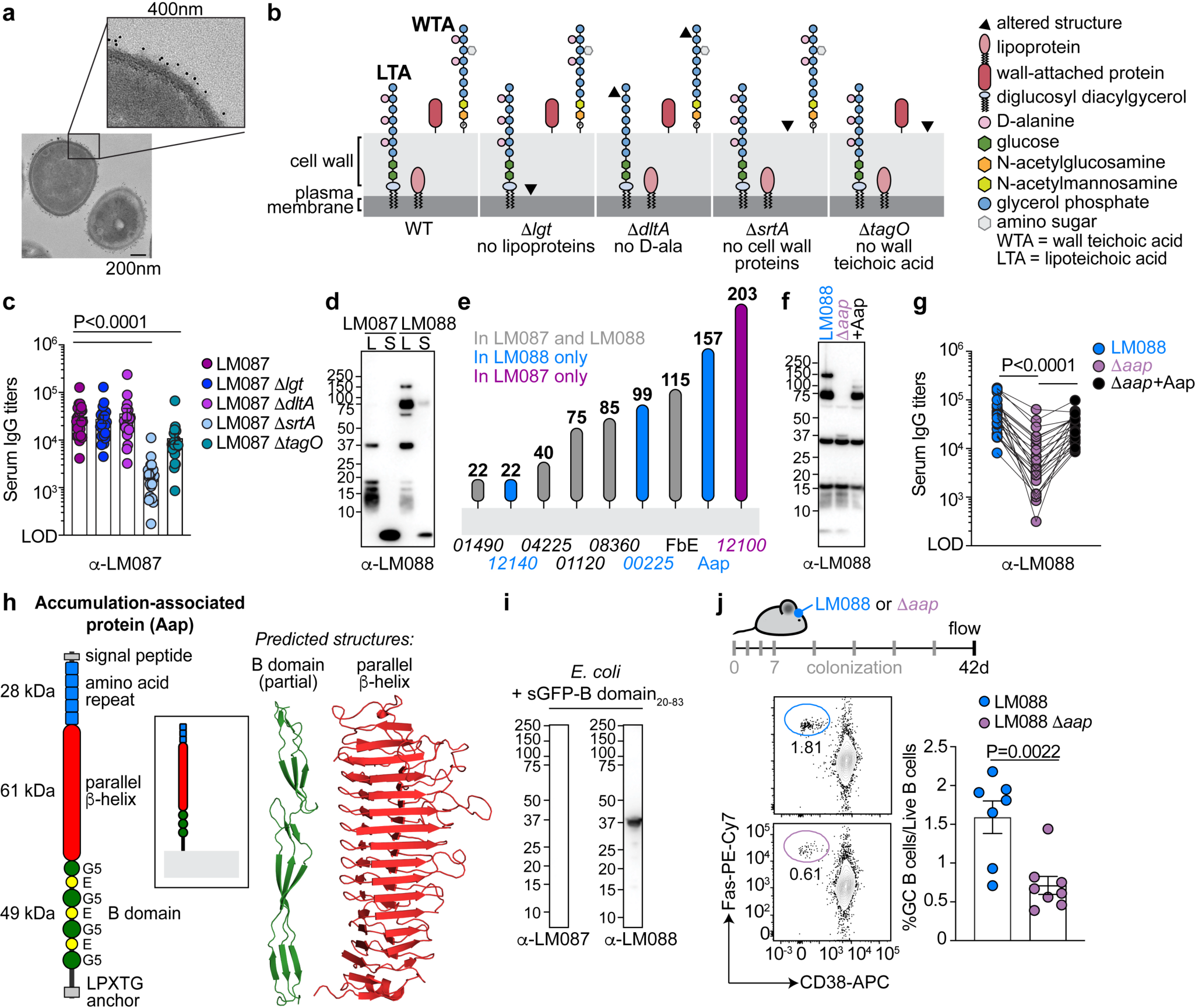
The predominant target of the antibody response is Aap. (**a**) Electron micrograph (and enhanced zoom) showing LM088 coated with IgG derived from the serum of mice colonized with LM088 for 6 weeks. (**b**) Schematic of the *S. epidermidis* cell surface mutants used in (**c**). (**c**) Bacterial ELISA using the serum of mice colonized with LM087 for 6 weeks against a panel of cell surface mutants (n=23, three independent experiments pooled). (**d**) Immunoblot analysis of cell lysate (L) and culture supernatant (S) from LM087 and LM088 using the serum of mice colonized with LM088 for 6 weeks. One representative of at least three independent experiments is shown. (**e**) Schematic showing predicted sortase substrates from the genomes of *S. epidermidis* LM087 (purple), LM088 (blue) or shared by both strains (grey). (**f**) Immunoblot analysis against LM088 (1, blue), LM088 Δ*app* (2, purple) or LM088 Δ*app* + plasmid-borne Aap (3, black) using the serum of mice colonized for 6 weeks with LM088. (**g**) ELISA of LM088 (blue), LM088 Δ*app* (purple) or LM088 Δ*app* expressing Aap recombinantly (black) using the serum of mice colonized with LM088 for 6 weeks (n=24, three independent experiments pooled). (**h**) Schematic of accumulation-associated protein (Aap) in LM088 (left) and predicted structure of its two main domains using AlphaFold2 (right). (**i**) Immunoblot against the lysate of *E. coli* expressing Superfolder-GFP (sGFP) fused to the minimal epitope identified in the B domain of Aap (aa 20-83) using the serum of mice colonized with LM087 or LM088 for 6 weeks. One representative of at least three independent experiments is shown. (**j**) SPF mice were colonized for 6 weeks with LM088 or LM088 Δ*app* for 6 weeks. The plot shows one representative mouse per group and the graph displays the percent of germinal center (GC) B cells among live B cells (Live^+^CD19^+^CD4^-^) in the cervical dLNs of all mice analyzed (n=7-8, two independent experiments pooled). Statistics: Graphs were analyzed using one-way ANOVA with Dunnett’s correction (**c**, **g**) for multiple comparisons and unpaired Student t tests (**j**). LOD: limit of detection. All graphs show mean +/− SEM.

To determine what class of surface molecule is bound by the antibodies, we used a set of LM087 mutants with defined cell envelope alterations^24^: Δ*lgt*, which is deficient in lipoproteins; Δ*dltA*, which is missing the D-alanine modification on teichoic acid; Δ*srtA*, which lacks cell-wall-attached proteins; and Δ*tagO*, which fails to elaborate wall teichoic acid (**Fig. 2b**). We immobilized each of these mutants to a plate and probed them with serum from LM087-colonized mice. Only Δ*srtA* showed a ∼10-fold reduction in antibody binding, indicating that a large fraction of the *S. epidermidis-*specific antibody pool binds to cell-wall-attached substrates of sortase (**Fig. 2c**). We obtained similar results by flow cytometry, confirming that the reduction in antibody binding to Δ*srtA* was not a trivial consequence of inefficient bacterial cell immobilization to the ELISA plate (**fig. s2a**). Though antibody binding to the Δ*tagO* mutant appeared partially reduced by ELISA, this defect was not observed by flow cytometry, suggesting that the apparent deficiency may have been caused by reduced binding of Δ*tagO* cells to the ELISA plate.

*srtA* encodes the enzyme sortase, which is a transpeptidase that recognizes a *C*-terminal sorting motif (LPXTG) in its substrates^25^. Catalytic attack by an active site cysteine on the amide bond between T and G yields an acyl-*S-*enzyme intermediate that is captured by the pentaglycine bridge on lipid II, linking a protein to a peptidoglycan precursor that will be incorporated into the cell wall. Having found that sortase is required for antibody recognition, we next sought to determine which sortase substrate contained the antigen. We used a computational approach to identify putative sortase substrates in *S. epidermidis*, searching the genomes of LM087 and LM088 for open reading frames that harbor an *N*-terminal signal sequence and an LPXTG motif within 50 residues of the *C*-terminus. We found six hits in LM087 and eight in LM088; of these, five are shared between the strains (**fig. s2b**).

We decided to focus on LM088, reasoning that it induces a stronger antibody response (see **Fig. 1a**). To determine which of its eight sortase substrates are bound by antibodies from murine serum, we ran a western blot against the cell lysate (L) and culture supernatant (S) of LM087 and LM088 using serum from LM088-colonized mice (α-LM088) (**Fig. 2d**); we used α-LM087 serum as a control (**fig. s2c**). A set of lower molecular weight bands were shared between LM087 and LM088. In contrast, we observed bands of approximately 150, 80, and 60 kDa that were unique to LM088 (**Fig. 2d**). We hypothesized that they corresponded to accumulation-associated protein (Aap), which has a molecular weight of 157 kDa, is processed proteolytically into smaller fragments^26^, and is the only LM088 protein in that size range with a sortase motif at its *C*-terminus (**Fig. 2e**).

To test this hypothesis, we constructed two derivatives of LM088: one in which *aap* is cleanly deleted (Δ*app*) and the other in which Δ*app* is complemented by a plasmid-borne copy of the gene (Δ*app*+Aap). A western blot against lysates from LM088, Δ*app*, and Δ*app*+Aap using serum from LM088-colonized mice revealed the disappearance of the prominent high-molecular-weight bands in Δ*app* and their reappearance in Δ*app*+Aap (**Fig. 2f**). Similar results were obtained by immobilizing these bacterial strains on a plate and probing them with serum from LM088-colonized mice (**Fig. 2g**). The titer of IgG that binds Δ*app* is 6-fold lower than wild-type, a deficit that is partially recovered by ectopic expression of Aap. These results indicate that Aap is a predominant (though not exclusive) target of the antibodies elicited by LM088.

In our initial experiments, we had observed the induction of mucosal IgA in the nares in response to colonization by wild-type LM088. We next asked whether IgA elicited by LM088 is also specific for Aap. We measured the quantity of IgA specific for LM088 and Δ*app* in the nasal washes of mice colonized with LM088 (**fig. s2d**). We observed a deficiency in binding of nasal IgA to LM088 Δ*app* starting at 6 weeks post-colonization, suggesting that the mucosal antibody response targets Aap, similarly to the serum antibody response. These results are consistent with a model in which Aap plays a role in systemic and local B cell responses to *S. epidermidis*.

Aap is a 1,431 amino acid protein composed of five domains: an *N-*terminal signal peptide; an amino-acid repeat domain (four complete and two partial 50-residue repeats); a large parallel β-helix domain of unknown function; a series of highly conserved G5-E repeat sequences (the B-domain) that form a fibril-like structure that extends upward from the cell wall^27^; and a *C-*terminal anchor domain that contains the sortase recognition sequence (**Fig. 2h**).

The Aap superfamily, which is widely distributed among human-associated *Staphylococcus* species^28^, includes proteins with different domains and domain organizations. For instance, the B domain—which is important for biofilm formation^29^—can contain up to 17 repeats, while the variant of Aap in LM088 contains only four. The best-known member of this family is present on the chromosome of *S. epidermidis* and contains an *N*-terminal lectin domain, which mediates binding to human corneocytes^30^. In contrast, the variant in LM088 is plasmid-borne and the lectin domain is replaced by a parallel β-helix domain of unknown function.

To identify the epitope(s) within Aap targeted by host antibodies, we constructed a series of truncation mutants and expressed the corresponding proteins recombinantly in *E. coli.* The parallel β-helix and B domains appear to be the primary epitopes, with the B domain bound most strongly (**fig. s3a-b**). Within the B domain, an additional set of truncated peptides narrowed the minimal epitope to a fragment of ∼60 amino acids but not further (**Fig. 2i** and **fig. s3c-f**), suggesting a conformational (rather than linear) epitope. This is consistent with the observation that the B domain repeats are unusually stable: they can refold even at high temperatures and under denaturing conditions^31^. On the one hand, recognition of the B domain is not surprising; it consists of alternating repeat sequences, and epitopes that are multivalent are known to induce strong B cell receptor (BCR) activation^32^. On the other hand, it is surprising to find that a protein so efficiently recognized by B cells is conserved on the surface of *S. epidermidis*, in a way that seems to invite an anti-commensal antibody response. This observation is reminiscent of the repeat sequences on the surface of *Plasmodium falciparum*^33^ and raises the question of whether the antibody response is somehow beneficial for *S. epidermidis*, as previously reported for the gut commensal *Bacteroides fragilis*^18^.

Finally, noting that LM088 induces higher antibody titers than LM087, we asked whether deletion of *aap* impacts the strength of the B cell response against LM088. We colonized mice with wild-type LM088 or Δ*app* for 6 weeks (**Fig. 2j**). Mice colonized by Δ*app* had a reduced frequency of germinal center B cells, suggesting that the presence of Aap drives a more vigorous B cell response to *S. epidermidis.* An immunoblot analysis confirmed that mice colonized with Δ*app* had no antibodies against Aap (**fig. s4**). Overall, these results indicate that Aap is a predominant antigen on the surface of *S. epidermidis* that helps determine the magnitude of the anti-commensal B cell response.

### Redirecting the B cell response against a non-native immunogen

In recent work, we showed that by colonizing mice with a variant of LM087 in which a non-native epitope is expressed on the surface, we could direct the anti-commensal T cell response against the non-native epitope^13^. In so doing, we elicited a robust antitumor immune response, constituting a novel form of vaccination.

We reasoned that our discovery of an antibody against Aap might enable a parallel strategy to redirect the B cell response against an unrelated pathogen. By colonizing mice with an engineered variant of *S. epidermidis* in which Aap is fused to a non-native immunogen, we hypothesized that it might be possible to elicit an antibody response against a target of interest (**Fig. 3a**).

**Figure 3:**
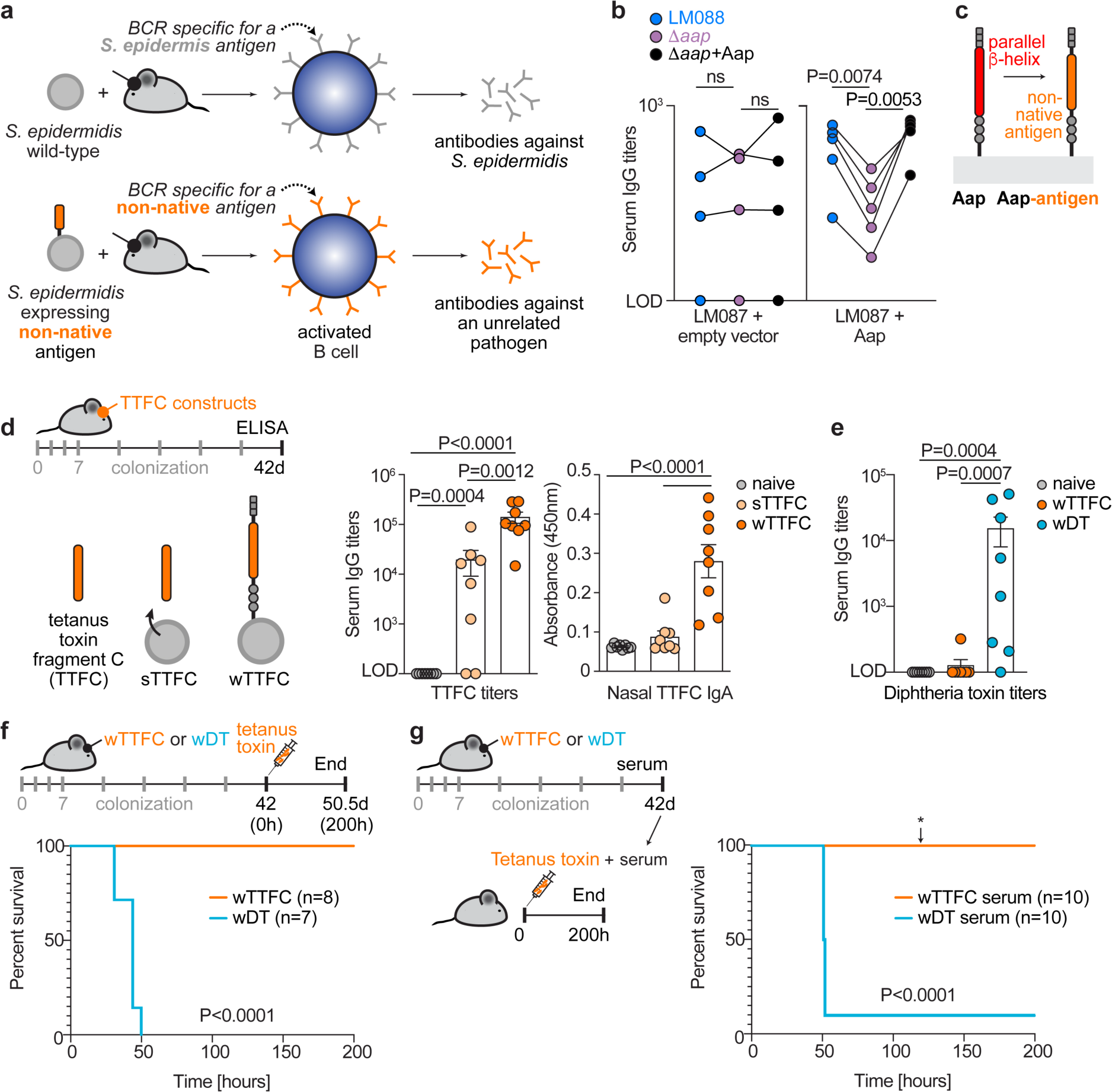
Redirecting the B cell response against a non-native immunogen. (**a**) Schematic of the approach used to redirect the B cell response toward a non-native antigen of interest. (**b**) Antibody titers against LM088, Δ*app*, and Δ*app* expressing plasmid-borne Aap (Δ*app*+Aap) in mice colonized for 5 weeks with LM087 harboring an empty vector (LM087 + empty vector) or expressing Aap on a plasmid (LM087+Aap). (**c**) Strategy used to express a non-native immunogen in *S. epidermidis*: an engineered version of Aap is expressed in which the parallel β-helix domain is replaced with the antigen. (**d**) SPF mice were colonized for 6 weeks with *S. epidermidis* LM088 Δ*app* engineered to express tetanus toxin fragment C (TTFC), a model immunogen. TTFC was either targeted for secretion (sTTFC) or attached to the cell-wall by fusion to Aap (wTTFC) and the quantity of TTFC-specific antibodies were measured in the serum (IgG serum titers) and in nasal washes (undiluted, absorbance at 450nm). (n=8/group, two independent experiments pooled). (**e**) IgG serum titers specific for diphtheria toxin (DT) in mice colonized with LM088 Δ*app* + Aap-TTFC (wTTFC) or + Aap fused to a catalytic mutant version of DT, Aap-DT (wDT) for 6 weeks. (n=8/group, two independent experiments pooled). (**f**) Mice colonized with either wTTFC or wDT were challenged with a lethal dose of tetanus toxin (150ng/kg) 6 weeks later. (n=7-8/group, two independent experiments pooled). (**g**) Mice were injected with a lethal dose of tetanus toxin (110ng/kg) preincubated with the serum of mice colonized with either wTTFC or wDT (serum shown in (**d**, **e**)). Of note, ∼120 h post injection, 4/10 mice in the wTTFC group develop mild tetanus symptoms which never reached the humane endpoint (see *). P values were calculated with one-way ANOVA with Dunnett’s (**b**, **e**) and Tukey (**d**) multiple comparison tests and log-rank tests (**f**, **g**). LOD: limit of detection. All graphs show mean +/− SEM.

To test this hypothesis, we started by engineering LM087 to express Aap (LM087-Aap), which is not present in its genome (see **Fig. 2e**). We colonized mice with LM087-Aap and then tested their serum antibodies for binding to wild-type LM088, Δ*app*, and Δ*app*+Aap (**Fig. 3b**). As a negative control, we colonized mice with LM087 harboring an empty vector. IgG from the mice colonized with LM087-Aap recognizes two Aap-expressing strains (LM088 and Δ*app*+Aap) but not an Aap-deficient mutant (Δ*app*), confirming a specific antibody response against Aap. In contrast, mice colonized with the negative control strain (LM087 + empty vector) did not have Aap-specific serum IgG. These data demonstrate that it is possible to elicit a B cell response against a non-native protein by expressing it on the surface of *S. epidermidis*.

Next, we introduced a B cell epitope from a pathogen of interest (**Fig. 3c**). As a model immunogen we chose tetanus toxin fragment C (TTFC), a 52 kDa *C*-terminal domain that is non-toxic^34^ but can elicit a protective antibody response against tetanus toxin^35^. Since we were uncertain of whether this would work—and if so, what the ideal scaffold would be to display the immunogen—we made chimeric versions of four carrier proteins bearing TTFC (**fig. s5 and s6**): the sortase substrates protein A (*Staphylococcus aureus,* 11 kDa)^13^, M6 (*Streptococcus pyogenes,* 27 kDa)^36^, and Aap; LysM (*S. epidermidis*, 18 kDa) which binds peptidoglycan non-covalently; and the signal sequence from FepB (*S. aureus*)^13^, targeting TTFC for secretion (sTTFC) into the culture medium.

To test whether these proteins elicit antibodies against TTFC in the setting of colonization, we engineered LM088 Δ*app* and LM087 to express each TTFC fusion protein. We then colonized mice for 6 weeks with these strains and measured the titer of TTFC-binding IgG in serum and IgA in nasal washes (**Fig. 3d**). Mice colonized with Aap-TTFC (wTTFC) had high titers of anti-TTFC IgG in serum and IgA in the nares, demonstrating that engineered *S. epidermidis* can elicit a robust systemic and mucosal antibody response against a non-native immunogen. Antibody titers elicited by the other chimeric proteins were lower or undetectable, indicating that the spatial and structural requirements for eliciting a B cell response are satisfied by Aap but not other carrier proteins (**fig. s6**). We note that protein A and the secreted antigen scaffold can elicit a productive T cell response against non-native epitopes^13^, suggesting that the antigen scaffolding requirements may be stricter for a B cell response than a T cell response.

Having seen that Aap is a productive scaffold, we made a series of variants of wTTFC (**fig. s5**) (i.e. smaller amino acid repeat domain, duplication of TTFC, or duplication of the B domain). While expressing the immunogen in place of the parallel β-helix domain gave rise to high antibody titers against TTFC (See **Fig. 3e**), other attempts to improve the carrier led to less robust B cell responses (**fig. s6**).

We next asked whether we could extend this strategy by replacing the parallel β-helix domain with a different immunogen (**Fig. 3e**). We cloned a catalytically dead mutant of diphtheria toxin (DT) into the same site, engineered this transgene into LM088 Δ*app*, and colonized mice topically with the resulting strain (wDT). After six weeks, wDT-colonized mice had high titers of IgG against DT in circulation, showing that this strategy is not limited to a single immunogen.

We reasoned that the antibody titers elicited by wTTFC were sufficiently high that they might be protective against a challenge with tetanus toxin^35^. We colonized mice for 6 weeks with wTTFC (or wDT as a negative control) and challenged them with a lethal dose of tetanus toxin. wTTFC-colonized mice were completely protected from death (**Fig. 3f**) and symptoms of toxin exposure (**Videos s1-2**), demonstrating that topical colonization with an engineered skin commensal can yield protection against a heterologous challenge. The mice colonized with wTTFC remained symptom-free for the duration of the experiment (**Video s3**).

Finally, we asked whether the protection was mediated by neutralizing antibodies. We isolated serum from colonized mice, incubated it with tetanus toxin, and then injected the toxin into mice (**Fig. 3g**). Pre-treatment with wTTFC serum (but not wDT serum) was sufficient to prevent death from toxin exposure, suggesting that the protective effect of wTTFC is mediated, at least in part, by antibodies. These experiments indicate that the antibody response generated by *S. epidermidis* colonization is sufficiently potent to provide protection against a lethal dose of a bacterial neurotoxin.

### Surface conjugation to expand the immunogen scope

Encouraged by these results, we sought to expand the scope of immunogens that can be displayed on the surface of *S. epidermidis*. We reasoned that the main limitation of the Aap chimera is that the immunogen needs to be functional when expressed in *S. epidermidis*—properly folded, secreted, and post-translationally modified if needed. Viral glycoproteins, which are large and heavily glycosylated, would be difficult to express in a functional form in bacteria.

To solve this problem, we devised an alternative approach that would enable us to express and purify a recombinant immunogen separately—e.g., in a mammalian cell—and then attach it to the surface of *S. epidermidis* (**Fig. 4a**). We took advantage of the SpyCatcher/SpyTag system^37^, which is derived from the extracellular *Streptococcus pyogenes* protein CnaB2, itself a sortase substrate. CnaB2 is stabilized by an autocatalytic isopeptide bond; by splitting this protein into two pieces, the globular SpyCatcher domain reacts quickly and irreversibly with the 16-aa SpyTag, forming a stable isopeptide bond (**Fig. 4b**).

**Figure 4:**
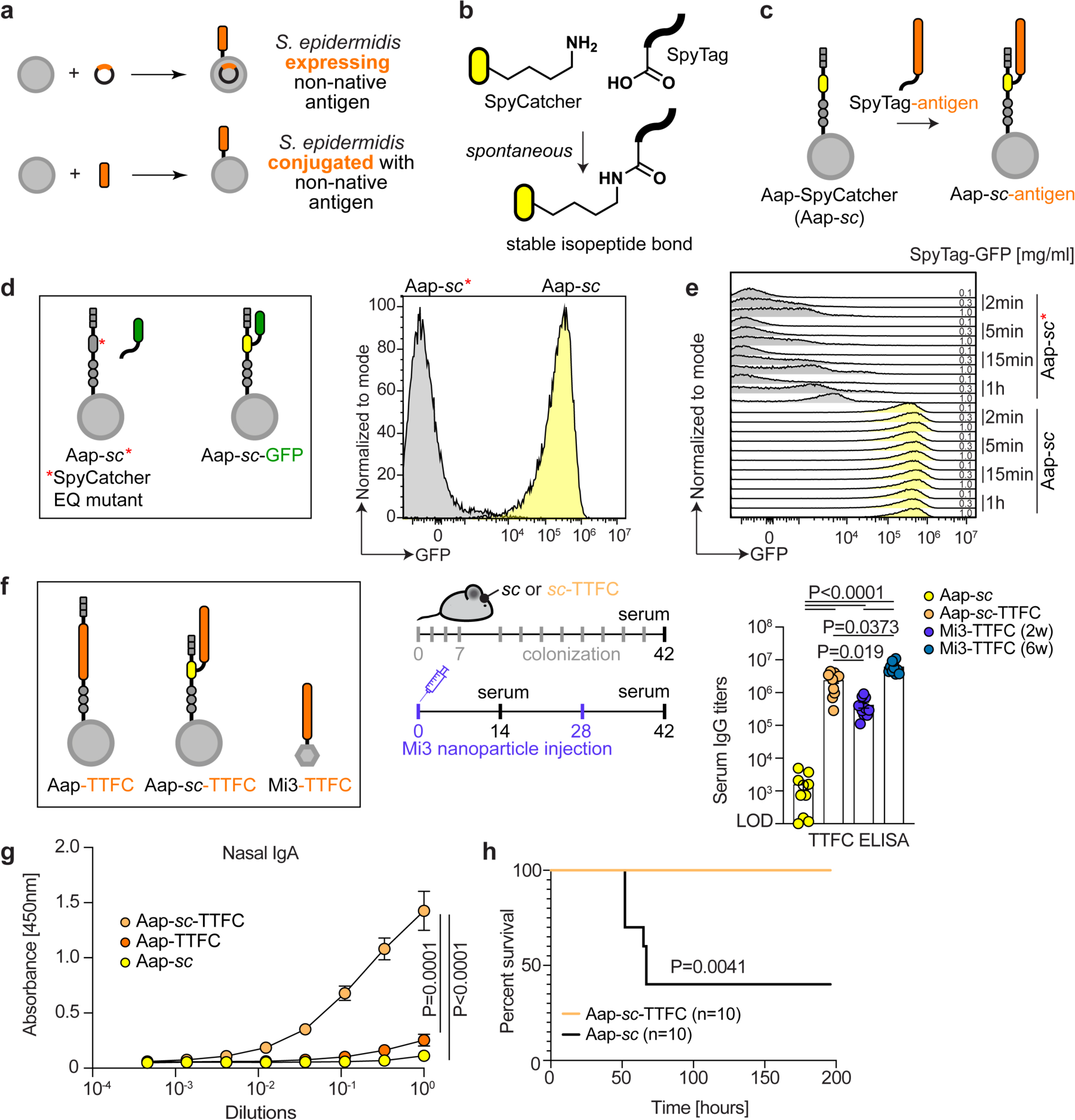
Generation of immunity using enzymatic conjugation. (**a**) Schematic representing the difference in approaches developed to elicit an antibody response against a non-native antigen using engineered *S. epidermidis*. (**b**) Chemical reaction between SpyCatcher (*sc*) and SpyTag-containing proteins. (**c**) Schematic representing the attachment of a non-native antigen to a strain expressing an Aap-SpyCatcher (Aap-*sc*) fusion. (**d**) *S. epidermidis* LM087 was engineered to express either a fusion between Aap and SpyCatcher (Aap-*sc*, yellow) or a catalytically dead version of SpyCatcher (Aap-*sc*\*, grey). Both strains were incubated with 0.1 mg/ml of sGFP-SpyTag for 2 min and visualized by flow cytometry. (**e**) Same as (**d**) but for 2, 5, 15 or 60 min and using 0.1, 0.3 or 1 mg/ml of SpyTag-GFP. (**f**) Mice were colonized with either Aap-*sc* conjugated with tetanus toxin fragment C-SpyTag003 (*sc*-TTFC) or the unconjugated strain (*sc*); antibody titers specific for TTFC were measured in the serum 6 weeks later. In parallel, another group of mice were immunized intramuscularly with SpyCatcher-based nanoparticles (mi3) conjugated with TTFC and antibody titers were measured in unboosted (2 weeks) or boosted mice (6 weeks). (**g**) TTFC-specific IgA in mice shown in (**f**) and in mice colonized with Aap-TTFC (wTTFC) expressing strain (mice shown in Fig. 3d). (**h**) Mice were injected with a lethal dose of tetanus toxin (110 ng/kg) preincubated with the serum of mice colonized with either *sc*-TTFC or *sc* (serum shown in (**f**)). P values were calculated with one-way ANOVA with Tukey’s (**f**) multiple comparison tests, two-way ANOVA with Dunnett’s post hoc test (**g**) and log-rank tests (**h**). Panels (**d**, **e**) are representative of three independent experiments and (**f**-**h**) show two independent experiments pooled. LOD: limit of detection. All graphs show mean +/− SEM.

We reasoned that a *S. epidermidis* strain expressing a chimeric version of Aap in which the parallel β-helix domain is replaced with SpyCatcher003 (a catalytically enhanced version of SpyCatcher^38^) could be conjugated with a recombinant protein bearing SpyTag003 (**Fig. 4c**). In principle, this could enable a wide range of recombinant proteins to be conjugated to the surface of *S. epidermidis*.

We expressed Aap-SpyCatcher003 in LM087 (hereafter, *sc*), incubated it with recombinant Superfolder GFP (sGFP) bearing a *C*-terminal SpyTag003 (**fig. s7a**), and analyzed bacterial cells by flow cytometry. After 2 min, *sc* cells are conjugated robustly with sGFP-SpyTag003, whereas a variant of this strain expressing a catalytically dead version of SpyCatcher (*sc*\*) is not efficiently conjugated (**Fig. 4d-e**). Thus, the *sc* strain enables highly efficient conjugation of recombinant proteins to the surface of *S. epidermidis*.

Next, we sought to determine whether a SpyCatcher-expressing strain conjugated with a recombinant immunogen could elicit an antibody response in vivo. We conjugated *sc* with TTFC-SpyTag003 (**fig. s7b**) and colonized mice with the immunogen-conjugated cells for six weeks (**Fig. 4f**). As a point of comparison, we prepared a nanoparticle vaccine—mi3-SpyCatcher conjugated with TTFC-SpyTag003, which contains up to 60 immunogens per particle^39^—and used it to immunize a second group of mice by intramuscular injection. To our surprise, serum from *sc*-TTFC-colonized mice had very high titers of anti-TTFC IgG (**Fig. 4f**): more than 10-fold higher than Aap-TTFC-colonized mice (**see Fig. 3e**). Remarkably, the response generated with *sc*-TTFC was of similar magnitude to the highly potent intramuscular mi3-TTFC vaccine. Mice colonized with *sc*-TTFC also harbored high concentrations of α-TTFC IgA in the nasal cavity, orders of magnitude higher than mice colonized with wTTFC (**Fig. 4g**).

Next, we asked whether this treatment protects mice against a lethal challenge with tetanus toxin. Mice challenged with tetanus toxin preincubated with *sc*-TTFC serum were fully protected against tetanus (**Fig. 4h**), confirming that *sc*-TTFC elicits potent neutralizing antibodies. Thus, display of antigens on the surface of *S. epidermidis* using the SpyCatcher system is an efficient strategy to elicit potent antibody response in vivo, and immunogen dilution over rounds of cell division does not result in weaker titers.

## DISCUSSION

Before this work, very little was known about B cell responses to skin colonists. Although some gut commensals induce an antibody response^6,18,40,41^, the titer of IgG induced by *S. epidermidis* is unusually high—comparable to what would result from a robust vaccination. The novelty and potency of the B cell response to *S. epidermidis,* combined with data from an accompanying manuscript shedding light on the mechanism of transdermal sampling by Langerhans cells^42^, reveals a powerful new facet of the program for immune surveillance of the microbiome.

The identification of Aap as a predominant target of the B cell response to *S. epidermidis* raises a number of questions. Aap is a large protein composed almost exclusively of sequence and structural repeats. It is tempting to speculate that these elements cross-link the B cell receptor, leading to strong B cell activation^43^ and the potent antibody response observed here. This is especially notable in light of the observation that the B domain—which contains the main epitope (**Fig. 2i** and **fig. s3c-f**)—is not only composed of repeats but also forms ordered aggregates^44^, creating even more crosslinking potential. The B domain is widely conserved among Gram-positive bacteria^45^ and is under selective pressure during an infection^46^, reminiscent of the conserved epitopes recognized by commensal specific CD4^+^ T cells in the intestine^5,11^ and suggesting that pre-emptive immune responses may protect against a broader set of strains than the elicitor. Immune recognition of repeat sequences in sortase substrates—in mice and humans^47^—may be more general than was previously known.

Our discovery of Aap as a predominant target of the antibody response enabled us to develop a method of topical vaccination. This approach differs in four ways from conventional vaccine modalities: (*i*) Protein-based subunit vaccines are more immunogenic when administered as multivalent antigens. Recent examples employ nanoparticle scaffolds for multivalent display (Gardasil, skycovione), facilitating B cell activation for a more robust antibody response^32^. *S. epidermidis* displaying an Aap-scaffolded epitope is, in essence, particulate immunogen with much higher valency than nanoparticle vaccines, and could be made even more so by increasing the number of immunogen copies in each Aap molecule. (*ii*) Most vaccines are administered by intramuscular injection. *S. epidermidis* is topical; it could be formulated in a cream and administered without a needle or a healthcare worker, which could facilitate low-cost distribution. (*iii*) A conventional vaccine is administered as a one-time bolus with subsequent booster doses. The biology of colonization is fundamentally different; antigen is encountered in a slow and steady fashion over the course of weeks. Conventional vaccines delivered with similar kinetics yield higher titer responses that target a more diverse set of epitopes^48^. (*iv*) Most traditional vaccines are administered by injection and result in an inflammatory response (i.e., fever and swelling). Vaccination with *S. epidermidis* via colonization does not require a barrier breach or induce inflammation.

Previous efforts to use bacterial strains to elicit an antibody response focused mainly on lactic acid bacteria^49–52^; antigen-specific responses have been observed both systemically and mucosally. However, it is unclear whether these examples involved sustained exposure at the native site of colonization and invoked a physiologic immune response to commensals. In contrast, we developed a commensal vaccine based on the natural response to *S. epidermidis* and its native surface protein Aap, across an intact skin barrier.

Our method has not yet been optimized. Given the antibody titers we are observing and the potency of protection against tetanus toxin challenge, it should be possible to administer a smaller number of doses without compromising prophylaxis. Moreover, we are only colonizing the head of the mouse; a broader surface area of colonization would involve a larger number of lymph nodes, presumably amplifying the magnitude of the response. There are numerous opportunities for improving the nature, localization, and copy number of the immunogen on the surface of the bacterial cell. Finally, using a combination of strains that redirect B and T cell responses could provide even broader immunologic protection against viral and bacterial pathogens.

It is particularly notable that colonizing the skin with *sc*-TTFC induces high titers of IgA in the nares. It is unclear whether this is due to incidental colonization of the nares, or instead involves the movement of IgA-producing B cells to the nares. If mucosal spreading is observed, it will be important to see whether there is an IgA response at other barrier sites, e.g., the lung and vaginal mucosa. This may also be of practical importance; mucosal vaccines hold great promise for preventing transmission by blocking infection at the site of pathogen entry, but they have been challenging to develop. A topical vaccine that induces a robust mucosal IgA response could be a low-cost solution to this problem.

The purpose of the immune response to colonists is coming into focus. Like the response to pathogens, it is potent and antigen-specific. Unlike the anti-pathogen program, the anti-commensal system is pre-emptive (it occurs in the absence of infection) and trans-barrier, involving active sampling between epithelial cells into the spaces in which colonists reside. In a very real sense, the anti-commensal program appears to vaccinate the host against the residents of certain barrier sites. Indeed, it may be that most of the encounters between the immune system and a microbe—and the majority of energy spent by the immune system—are devoted to this purpose.

## ACKNOWLEDGMENTS

We are deeply indebted to members of the Fischbach lab for helpful discussions and suggestions, especially Brian Caliando for cloning support, Martin McLaughlin for chemistry questions and thorough reading of the manuscript, and Alice Cheng for microbiology input; members of the Victora lab, especially Gabriel Victora, Ariën Schiepers and Luka Mesin for sharing protocols and insightful conversations; Angelina Bilate, Julie Segre, Sean Conlan, and Heidi Kong for providing feedback and helpful discussions; Maria Prado, Amanda Espinoza and Joanne Au for keeping the lab running; Marco Pirazzini and Cesare Montecucco for assistance with the tetanus toxin challenge experiments; members of the Stanford University Veterinary Service Center for animal husbandry; the electron microscopy facility of Washington University, especially Gregory Strout for guidance with EM sample preparation; and staff members of the Stanford University shared FACS facility for assistance with flow cytometry analysis (NIH grant 1S10OD026831-01). This work was supported by the Stanford Microbiome Therapies Initiative, the Swiss National Science Foundation (D.B., Early Postdoc.Mobility and Postdoc.Mobility), NIH grants 5R01AI175642-02 (M.A.F.) and 1F32HL170591-01 (L.J.B.), the Howard Hughes Medical Institute (Y.E.C., C.O.B., Hanna H. Gray Fellowship), the Bill and Melinda Gates Foundation (M.A.F.), Open Philanthropy (M.A.F.), the Leona M. and Harry B. Helmsley Charitable Trust (M.A.F.), the Chan Zuckerberg Biohub (C.O.B., M.A.F.), the Division of Intramural Research of the National Institute of Allergy and Infectious Diseases (NIAID) (Y.B. and I.G.), the Office of Research Infrastructure Program, Office of The Director, National Institutes of Health under Award Number P51OD011107 (CNPRC, UC-Davis), the Department of Defense NDSEG Fellowship (P.V.L.) and the Knight-Hennessy Fellowship (P.V.L.).

## AUTHOR CONTRIBUTIONS

Conceptualization: D.B., K.D.B., Y.E.C., Y.B., C.O.B., M.A.F., Methodology: D.B., K.D.B., Y.E.C., I.G., Y.B., C.O.B., M.A.F., Investigation: D.B., K.D.B., Y.E.C., V.K.Y., P.V.L., A.N., A.V., J.L.P., T.T.D.N., J.M.S., Y.E.L., A.D., J.L.P., S.J., X.M., T.T.D.P., A.Z., L.J.B., Visualization: D.B. and M.A.F., Funding acquisition: D.B., Y.E.C., P.V.L., L.J.B., K.K.A.V.R., Y.B., C.O.B., M.A.F., Supervision: D.B., K.D.B., Y.E.C., K.K.A.V.R., Y.B., C.O.B., M.A.F., Writing – original draft: D.B. and M.A.F., Writing – review editing: D.B., K.D.B., Y.E.C., V.K.Y., P.V.L., L.J.B., I.G., K.K.A.V.R., C.O.B., M.A.F..

## COMPETING INTERESTS

M.A.F. is a co-founder of Kelonia and Revolution Medicines, a member of the scientific advisory boards of the Chan Zuckerberg Initiative, NGM Biopharmaceuticals, and TCG Labs/Soleil Labs, and an innovation partner at The Column Group. D.B., Y.E.C., K.D.B., C.O.B, Y.B. and M.A.F. are inventors on patent applications submitted by Stanford University and the Chan Zuckerberg Biohub that cover methods for using engineered bacteria to elicit antigen-specific immune cells.

## SUPPLEMENTARY MATERIALS

Materials and Methods

Figures s1-s7

Videos s1-s3

Table s1

