## Supplement for "Discovery and engineering of the antibody response against a prominent skin commensal"

**This PDF file includes:**

Materials and Methods
Figs. s1 to s7
Videos s1-3

### MATERIALS AND METHODS

#### Mice

Mice were housed at Stanford University under specific pathogen-free (SPF) conditions according to the guidelines established by the Stanford University institutional animal care and use committee. C57BL/6 (B6) and *rag2*$^{-/-}$ (RAGN12) mice were purchased from Taconic.

#### Skin colonization

6-10-week-old C57BL/6 mice were colonized with the indicated *Staphylococcus epidermidis* strains similarly to a previously described protocol[1]. Briefly, overnight cultures of *S. epidermidis* were normalized to an $OD_{600}$=6 with BHI (or BHI + chloramphenicol (10 µg/ml) where appropriate, Sigma). Normalized bacterial cultures were gently applied on the head, ears and nose of mice with sterile cotton swabs (Puritan, 25-8062PD). The colonization was repeated every other day for one week for a total of four associations and then once/week until the animals were sacrificed unless otherwise indicated on the figure or the legend. For serum harvest, we typically harvested blood at the experimental endpoint (to avoid inserting bacteria into the blood stream) by cardiac puncture. The nasal washes were harvested using 1ml of PBS (1ml syringe attached to a 22G catheter, Exel International, 26746). Briefly, the catheter was inserted upward into the trachea and the nasal cavity was gently flushed and the nasal wash harvested in an Eppendorf tube at the entrance of the nose.

#### Bacterial cultures

*S. epidermidis* was grown on Difco Brain Heart Infusion (BHI) agar plates (BD 241810) overnight at 37ºC. Single colonies were picked, inoculated into BHI (Difco Brain Heart Infusion media, BD 237200), and grown overnight at 37ºC with shaking (220 rpm). When the strain carried a pLI50-based plasmid, the bacteria were grown on/in BHI + chloramphenicol (10 µg/ml) (Sigma, C1919).

#### NHP skin bacterial isolation

Non-human primate (NHP) *S. epidermidis* strains were isolated from the skin of macaques similarly to a previously described method[2]. All macaques were healthy adult rhesus macaques (Macaca mulatta), 10-18 years of age, housed at the California National Primate Research Center (University of California, Davis), which is accredited by the Association for Assessment and Accreditation of Laboratory Animal Care International (AAALAC). Animal care was performed in compliance with the 2011 Guide for the Care and Use of Laboratory Animals provided by the Institute for Laboratory Animal Research. Briefly, the unprepped skin (elbow and abdomen) of ten macaques was gently swabbed with ESwabs (Thermo Fischer, R723481) (one swab/monkey). The swabs were then returned to the LoBind Eppendorf tubes containing buffer from the kit (according to manufacturer) and shipped to Stanford for microbial isolation.

To avoid cross-contamination, new gloves were used between each monkey. The next day, the sample solution was grown on BHI plates and grown aerobically overnight. Single bacterial colonies from the original plates were grown on fresh BHI plates, grown overnight aerobically, identified by MALDI-TOF[3], isolated, stocked, and stored at -80ºC. For each sampled monkey, we also collected serum to test antibody titers against the isolated strains.

**Flow cytometry of mammalian cells**

Cervical lymph nodes were harvested from mice, smashed on a 100 µm cell strainer (Miltenyi Biotec, 130-110-917) using the back of a 1 ml syringe (Becton Dickinson, 309659) in filter sterilized (MILLIPORE S2GPU05RE) PBE buffer (Miltenyi Biotec, one bottle of autoMACS Rinsing Solution supplemented with one bottle of BSA stock solution, 130-091-222 and 130-091-376). Dissociated cells were stained for surface markers at 4°C for 30 min in PBE. When cells could not be run on the same day, the cells were fixed in either BD cytofix/cytoperm or using the eBioscience Foxp3/Transcription Factor Staining Buffer Set (Invitrogen) according to manufacturer instructions. Flow cytometry data was acquired on a FACSymphony or a FACS LSR II machine (Becton Dickinson) and analyzed using FlowJo software package (Tree-Star).

**Bacterial ELISA**

High-binding ELISA plates (Corning, 9018) were precoated with 100 µl of poly-L-lysine solution (Sigma-Aldrich, P4707-50ML) for 15 min at room temperature. The plates were then washed three times with 200 µl of PBS, dried and coated with bacteria. The bacterial cultures were prepared as follows: overnight cultures of the indicated bacteria were washed two times with PBS (Thermo Fisher Scientific, 10010049) and normalized to an $OD_{600}$=6 in PBS. 50 µl of normalized cultures were used to coat each well. The plates were sealed (Bio-Rad, MSB1001) and incubated overnight at 4°C. The following day, the plates were washed five times with PBST (PBS + 0.05% Tween-20, Sigma, P1379), blocked with 200 µl of 2% BSA (Sigma-Aldrich, A7030-100G) in PBST at room temperature for 2 h. The plates were then incubated with the indicated sera from mice in serial dilutions (typically 1:100 first dilution and then 1:3 or 1:4). For nasal wash ELISA, we used undiluted samples and then performed 1:3 dilutions. Next, the plates were washed five times with PBST and incubated with secondary antibody at a dilution of 1:10,000 (mouse IgG: SouthernBiotech, 1030-05; Mouse IgA: SouthernBiotech, 1040-05, Human IgG: Thermo Fischer, 31413; NHP IgG: Thermo Scientific, PA1-84631, see supplementary details for other secondary antibodies) in 2% BSA/PBST. After five more PBST washes, the plates were revealed using 100 µl of TMB reagents (BD, 555214) and the reaction stopped with 50 µl stop solution (BioLegend, 423001). Antibody titers were calculated by logarithmic interpolation of the dilutions with absorbance at 450nm immediately above and below the absolute value 0.2, as previously described[4].

**Bacterial flow analysis**

Overnight cultures of bacteria were washed 2x with PBS and normalized to 1 ml of $OD_{600}$=1. The bacteria were then centrifuged (12,000 x *g* for 5 min at 4°C) and resuspended in 50 µl of blocking buffer (PBS containing 5% serum from *Rag2*$^{-/-}$ mice) and incubated for 45 min at 4°C. The bacteria were pelleted and then resuspended in 100 µl of PBS containing 2% serum from mice colonized with *S. epidermidis* for the indicated times, and incubated for 2 h at 4°C. The bacteria were then washed 2x with 1 ml of PBS, resuspended in 50 µl of $\alpha$-mouse IgG-PE-Cy7 (1:100), and incubated for 40 min at 4°C. Next, 1 ml of Syto9 solution (1.5 µl Syto9/1 ml PBS) was added and incubated for 15 min at RT. The bacteria were then spun down, washed 2x with PBS, resuspended in 200 µl of PBS, and analyzed by flow cytometry. Flow cytometry data was acquired on a FACSymphony (Becton Dickinson) or a CytoFLEX machine (Beckman Coulter) and data were processed using the FlowJo software package (Tree-Star).

**Immunoblotting analysis**

For *S. epidermidis* immunoblotting: overnight cultures were normalized to an equivalent of 500 µl of $OD_{600}$=8, washed 2x with 1 ml of TBS (50 mM Tris-HCl, pH 7.5, 150 mM NaCl), resuspended into 100 µl of TBS containing 100 µg/ml of lysostaphin (Sigma, L7386) from 50x stocks (5 mg/ml) and incubated for 2 h at 37ºC. Of note, for the immunoblots shown on panels on Fig. 2d and fig. s4, bacteria were lysed in TBS + 100µg/ml lysostaphin + 20% sucrose. The lysates were then incubated on ice for 5 min; 1:500 benzonase (Sigma, E8263) was added and samples were incubated on ice for 15 min or longer to ensure reduced viscosity (digestion of DNA). For supernatants, after centrifuging overnight cultures, the supernatant was concentrated using an Amicon 3k column (UFC500396) at 14,000 x *g* at 4°C for 20 min. The samples were then prepared for SDS-PAGE: normalized to protein content (0.2 mg/ml final) and supplemented with 100 µM DTT and loading buffer (Life Technologies, NP0007) according to manufacturer recommendations. The samples were heated at 95ºC for 5 min, loaded on an SDS-PAGE gel (Thermo Scientific, NP0336BOX or NP0335BOX), and run in MES buffer (Thermo Scientific, NP0002) at 150 V for ~2 h at 4ºC. The proteins were transferred onto PVDF membranes using the Trans-Blot Turbo Transfer System (Biorad). Membranes were blocked for 1 h at RT in 5% milk in TBS-T (TBS + 0.1% Tween-20, Sigma, P1379) and incubated with primary antibody (1:500 for serum, 1:2,000 for anti-HA-HRP) for 1 h at RT in 5% milk in TBS-T, washed 3x with TBS-T and incubated with secondary antibody ($\alpha$-mouse IgG, 1:10,000, Fisher Scientific, 62-652-0, no secondary for HA-HRP) for 1 h at RT in 5% milk in TBS-T, washed 3 more times, developed with SuperSignal reagent (Thermo Scientific, 34577), and imaged on an Azure Biosystems C200.

**Dot blot analysis**

Peptides were ordered with *N*-terminal biotinylation from Genscript with a purity of at least 85%. The peptides were resuspended in Mill-Q (MQ) water (EMD Millipore) and diluted to 1 mg/ml, 0.1 mg/ml and 0.01 mg/ml. On a nitrocellulose membrane (BioRad, 1620112), 2 μl of peptide (from the dilutions above) was loaded (see figure and legend for organization) corresponding to 2 μg, 0.2 μg and/or 0.02 μg peptide per dot. As positive controls, 2 μl of LM088 lysate (~20mg/ml) and 2 μl of mouse IgG1 isotype control (Biolegend, 400102, 0.5mg/ml) were also loaded on the blot. The membrane was air-dried for 30 min at RT and blocked for 1 h with 5% milk/TBST (see immunoblot for details). The membrane was then washed once with TBST and incubated with either a primary antibody (1:500 of serum of LM088-colonized mice for 6 weeks) or streptavidin-HRP (ThermoFisher, 21130, to ensure proper loading of the peptides) in 5% milk/TBST overnight at 4°C. The membrane was then washed three times and incubated with anti-mouse IgG-HRP secondary antibody (α-mouse IgG, 1:10,000, Fisher Scientific, 62-652-0) for 1 h at RT, washed 3 more times with TBST, developed with SuperSignal reagent (Thermo Scientific, 34577), and imaged on an Azure Biosystems C200.

**Immunogold electron microscopy**

Overnight cultures of *S. epidermidis* LM088 were normalized to 1 ml of $OD_{600}$=4, pelleted, washed 1x with 40 ml of PBS, and blocked in 600 μl of PBS/BSA 5% for 4.5 h at 4°C with rotation. Then 1.2 ul of primary antibody (1:500 dilution) was added and incubated with rotation overnight at 4°C. For the primary antibody, we used the serum of mice colonized with LM088 for 6 weeks and boosted once/week. The next morning, the bacteria were washed 3x with 1 ml of PBS and incubated with secondary antibody conjugated to colloidal gold (1:20 in 200ml/sample) for 4 h at 4°C in PBS/BSA 5% (6 nm Colloidal Gold AffiniPure Goat Anti-Mouse IgG (H+L) (EM Grade), Jackson Immunoresearch, 115-195-166). The bacteria were then washed 3x with 1 ml of PBS and fixed in 1 ml of 2.5% glutaraldehyde + 2% PFA in 0.1 M cacodylate buffer (pH 7.2) (Electron Microscopy Sciences, 16220, 11652 and 15710-S) for 1h at RT (rocking). Fixed samples were shipped to the Center for Cellular Imaging of Washington University for downstream processing and imaging.

***E. coli* competent cells**

Chemically competent *Escherichia coli* (*E. coli*) DC10B were prepared as follows: single colonies of DC10B from LB agar plates (BD, 244510) were inoculated into LB broth (Thomas Scientific, 244610) and grown overnight at 37ºC with shaking (220 rpm). The next day, the cultures were diluted 1:200 in LB and grown to $OD_{600}$~0.4. The bacteria were then rested on ice for 10min and centrifuged at 1,600 x *g* for 9 min at 4ºC and resuspended in 0.2x volume (e.g., for 100 ml of culture, resuspend into 20ml) of chilled $CaCl_2$ mix (for 1L: 120 ml 0.5M $CaCl_2$, 375 ml 40% w:v glycerol, 3.02 g PIPES, MQ to ~900 ml, adjust pH to 7.0 with NaOH and add MQ to 1L). The bacteria were then centrifuged for 7 min at 1,100 x *g* at 4°C and resuspended in 0.2x volume of chilled $CaCl_2$ mix and incubated on ice for 30 min. Next, the bacteria were centrifuged for 7 min at 1,100 x *g* at 4°C and resuspended into 0.04x volume of chilled $CaCl_2$ and incubated for 1h on ice. The bacteria were then aliquoted, flash frozen and stored at -80ºC.

**Cloning**

For all constructs, the inserts and backbones were amplified using PrimeStar MAX DNA Polymerase (Takara Bio, R045B) and purified using either a PCR cleanup kit (Zymo, D4014) or gel extracted if more than one band was visible on a diagnostic agarose gel. When a plasmid was used as template for the PCR reaction, an additional Dpn1 (NEB, R0176L) digestion was performed prior to purification, following manufacturer's instructions. When a colony was used as template, we followed the colony PCR protocol described below. Gel extraction was performed as follows: a band containing the desired DNA amplicon was excised from an 0.8-1% agarose gel and incubated with 500 μl of Qiagen QG buffer (Qiagen, 19063) at 95ºC for 5 min or for an additional 2 min if the gel was not completely melted. Next, up to 600 μl of the solution was added to a Zymo-Spin IC Columns (Zymo, C1004) centrifuged at max speed for 30 sec on a tabletop centrifuge. This step was repeated until all the solution was loaded onto the column. After the last spin, the column was washed 2x with 500 μl of PE buffer (Qiagen, 19065), spun at max speed, and eluted with 16 μl of nuclease-free water pre-heated to 95ºC (wait 1-2 min and centrifuge at 8,500 x *g* for 30 sec). The eluate was reloaded onto the same column to increase the yield of DNA recovered. To build the vectors, the purified PCR products were assembled using Gibson assembly (NEB, E2621L). For pLI50- and pIMAY-based vectors, the Gibson mixes were transformed into chemically competent DC10B (see below for protocol). Briefly, 1/3 of a Gibson reaction was added to 100 μl of chemically competent DC10B cells, and incubated on ice for 30 min, heat-shocked at 42°C for 45 sec, recovered at 37°C in LB for 1 h and then plated on the appropriate LB plate (LB+ Carbenicillin 50=50ug/ml (Teknova, C2105) for pLI50 and LB+chlor10 for pIMAY) for overnight growth. Proper assembly was confirmed by colony PCR and sequencing using OneTaq Quick-Load (NEB, M0486L). Correct clones were inoculated into liquid cultures (LB+ Carbenicillin 50=50ug/ml (Teknova, C2105) for pLI50 and LB+chlor10 for pIMAY) for overnight cultures, the plasmids miniprepped (Qiagen, 27104) and sent for sequencing. For pET-30b vectors, assembled Gibson mixes were transformed into stable competent cells (NEB, C3040H), the vectors confirmed by sequencing (see above) and eventually transformed into competent BL21 (DE3) bacteria for protein expression (NEB, C2527H). For pET-30b vectors, bacteria are grown in/on LB + Kanamycin50 = 50 ug/ml (VWR, AAJ61272-09).

**Construct design**

All antigen vectors for expression in *S. epidermidis* were designed similarly to what we previously published[5] (see fig. s5 and supplementary table s1 for sequences and more details). Briefly, all vectors were built on a replicative pLI50 plasmid backbone containing a constitutive promoter (Ppen) and a ribosome binding site from *Staphylococcus aureus* delta-hemolysin (hld) gene. Unless otherwise noted, all sequences were codon optimized for *S. aureus* using the IDT codon optimization tool. For repeated protein sequences, we used a simple algorithm to generate codon optimized DNA sequences that are distinct from each other to avoid recombination. Constructs where the antigen was fused to protein A fragment were based on a previous publication[5]. Briefly, the wall-spanning region of protein A XM domain was fused at the *C*-terminus to the antigens. The signal sequence from protein A was used as described[5]. For secreted constructs, we used the *N*-terminal Tat secretion signal from *S. aureus* fepB (gene locus name SAOUHSC_00326)[5] fused to the antigen, with no carrier protein. To build fusions to the peptidoglycan binding protein LysM, we fused the LysM protein from LM088 (*N*-terminus) to a fragment of LysM from *S. epidermidis* strain RP62A followed by the antigen domain. To build Aap-TTFC 2x B domain, we replaced the B domain from *S. epidermidis* LM088 on the original Aap-TTFC construct with the B domain of *S. aureus* COL strain, which is roughly twice as large. For plasmids for expression in *E. coli*, all constructs were cloned into a pET-30b vector, similarly to a previously described strategy[6]. To map the epitope within the B domain of Aap, we cloned truncations fused to a superfolder GFP (sGFP) protein to visualize the fragments by immunoblotting. To generate pIMAY-Aap, ~1 kb upstream and ~1 kb downstream of Aap were cloned into pIMAY vector, similar to a previous method[7]. The cloning strategy for all constructs, including all primers and gblock sequences, is available in Supplementary Table S1.

**Transformation of *S. epidermidis***

Genetic engineering of *S. epidermidis* was performed as previously described with minimal changes[5]. Briefly, *S. epidermidis* was grown overnight in BHI (Difco Brain Heart Infusion media, BD 237200) containing 0.5 M sorbitol (BHIS) (Sigma, S6021) at 37ºC shaking (220 rpm). The next day, the cultures were diluted 1:200, grown to $OD_{600}$ ~0.5-0.8, centrifuged at 3,600 x *g* at 4ºC, washed 5x with ice cold 10% glycerol using the same volume as the volume of cultures (i.e. for 40ml of culture, wash 5x with 40ml of 10% glycerol), and resuspended in 10% glycerol (for every 40 ml, use 70 μl of 10% glycerol =~ 2 transformations). During the washes, the DNA (previously isolated from *E. coli* DC10B strain) was desalted on 0.025 μm VSWP membrane filters (Sigma, VSWP02500). Briefly, 22.5 ml of MQ was added to a petri dish and one membrane was floated on the water. DNA was carefully added to the membrane for dialysis for 1 h at RT. Next, 100 μl of resuspended bacteria (see above) was added to Eppendorf tubes containing 0.2-1 μg of desalted DNA, gently mixed and left on ice for ~5 min. For the transformation, the bacteria and DNA were heat shocked for 2 min at 56ºC and immediately transferred to a 0.1 cm cuvette (Bio-Rad,

1652089) and electroporated with a pulse of 1.8 kV using a Bio-Rad Micropulser. The typical constant time was 2.3-2.5 and anything below 2.2 generally did not yield any transformants. After the transformation, the bacteria were transferred into 3 ml of BHIS and moved to a 37ºC shaker. After ~1h of recovery, the bacteria were plated on BHIS+chlor10 plates and grown overnight at 37ºC. The next day, transformants were grown on BHI+chlor10 plates and confirmed by colony PCR. Single colonies of correct clones were picked and inoculated in BHI + chlor10 for overnight cultures with shaking at 37ºC. The next day, the strains were banked, and the DNA sequence confirmed again by either miniprep (for LM087) or PCR (LM088) to ensure no mutations were acquired during liquid growth.

***S. epidermidis* colony PCR**

Single colonies of *S. epidermidis* were picked and added to 10μl of DMSO (Tocris, 3176), heat killed at 95°C for 30 min, and mixed with 20 μl of nuclease-free water (NEB, B1500S). 2.5 μl of the mixture was used in 25 μl PCR reaction using PrimeStar MAX DNA Polymerase (Takara Bio, R045B) following manufacturer's guidelines. PCR amplicons were run on a diagnostic agarose gel to confirm size and send for sequencing. For pLI50-based plasmids, primers DB394 + DB397 were used and for pIMAY DB212 + DB213.

***S. epidermidis* minipreps**

Single colonies of *S. epidermidis* were inoculated in 4 ml of BHI+chlor10 and grown overnight at 37°C with shaking. The next day, the cultures were centrifuged at 3,600 x *g* for 8 min, resuspended in 700 μl of TE buffer (10 mM Tris-HCl, pH 8.0, 1 mM EDTA) and transferred to an Eppendorf tube. Next, 500 μl of ice-cold acetone mix (50% acetone, 50% absolute ETOH, stored at -20°C) was added and incubated for 5 min on ice. The bacteria were then centrifuged for 2 min at 14,000 x *g*, washed 1x with 1 ml of TE buffer and the pellet resuspended in 250 μl of P1 buffer (Qiagen, 19051) supplemented with 3 μl of lysostaphin (Sigma, L7386) and incubated for 1-2 h at 37°C without shaking. Following this step, a normal miniprep procedure was performed (Qiagen, 27104). Of note, this procedure is not used for strain LM088, which contains its own native plasmids, making sequencing of the transformed plasmid more challenging.

**Generation of Aap-deficient *S. epidermidis* strain**

Generation of an Aap-deficient strain of *S. epidermidis* LM088 was performed as previously described[7] and as follows: *S. epidermidis* LM088 was transformed with pIMAY-Δ*aap*, recovered at 28°C for 4 h with shaking and plated on BHIS+chlor10 plates and grown for 2 d at 28°C. Single colonies from the electroporation plates were grown on BHI+chlor10 and grown for another 2 d at 28°C. At this point, we confirmed that the plasmid was intact and did not mutate by sequencing two amplicons: the temperature sensitive repA gene (DB219 + DB220) and the region flanking the Aap gene (DB212 + DB213). Once the sequences were confirmed, single colonies were inoculated into BHI+Chlor10 and grown with shaking at 37°C for ~8 h to cure the plasmid and then plated onto BHI+Chlor10 plates and grown overnight at 37°C. Single colonies were picked and grown on BHI+Chlor10 overnight at 37°C. At this step, we performed colony PCR and confirmed that the plasmid was cured (i.e, single crossover integration in the genome) using primers DB212 + DB213. Colonies that had successfully lost their plasmids were grown in BHI at 37°C for 6 h and plated onto BHI + anhydrotetracycline (1 μg/ml) to select for double cross-overs and grown for 2 d at 28°C. At this stage, single colonies were patched onto BHI + anhydrotetracycline and BHI+Chlor10, grown at 37°C overnight. The next day, single colonies that had grown on BHI + anhydrotetracycline but not on BHI+Chlor10 were screened for the intended KO by colony PCR using primers outside the homology arms, DB280 + DB281.

**SpyCatcher conjugation for in vitro experiments**

Overnight cultures of *S. epidermidis* LM087 + Aap-SpyCatcher003 (*sc*) or the catalytic mutant (*sc**, E77Q mutant of SpyCatcher) were OD normalized (150 μl of $OD_{600}$=8/sample), washed 2x with 1 ml of RT PBS and incubated for the indicated times (2, 5, 15, or 60 min) with the indicated amount of sGFP-SpyTag003 (0.1, 0.3 and 1 mg/ml) in PBS at RT (total volume=150 μl/sample). The cells were washed with 1 ml of PBS, spun down and resuspended in 100 μl of Mycolight staining buffer (1:100 dilution in PBS, MycoLight™ Red JJ94, AAT Bioquest) and incubated for 15 min at RT. The cells were then washed two times with 1 ml of PBS and resuspended in 200 μl PBS for flow cytometry analysis.

**SpyCatcher conjugation for in vivo colonizations**

Overnight cultures of *S. epidermidis* strain *sc* were OD normalized (5 ml/cage of $OD_{600}$=6), washed 2x with 50 ml of PBS and incubated for 15 min with 0.1 mg/ml of TTFC-SpyTag003 in PBS at room temperature (total volume=5ml/cage). The cells were washed with 50 ml of PBS 3x and resuspended in 5 ml/cage of BHI + chloramphenicol (10 μg/ml). The mice were then colonized as described above.

**Protein production and purification**

Superfolder-GFP (sGFP)-SpyTag003 and TTFC-SpyTag003 were cloned into a pET-30b expression vector. All expression vectors were transformed into BL21 (DE3) strain (NEB, C2527H) for protein purification. Single colonies of each construct were inoculated into LB with kanamycin (50 μg/ml) for overnight growth at 37°C with shaking at 220 rpm. The next day, the cultures were diluted 1:200 into 1L of Terrific Broth (Fisher Scientific, AAJ75856A1) + Kanamycin (50 μg/ml). When the cultures reached an $OD_{600}$=~0.4-0.6, protein expression was induced by adding 1 mM IPTG and transferring the cultures to 28°C with shaking at 250 rpm. The next day, the cultures were centrifuged (3,600 x *g* for 8 min at 4°C) and resuspended in prechilled 30 ml lysis buffer (50mM Tris-HCl pH 7.5 + 150 mM NaCl + 10 mM imidazole) plus DNAse (Sigma, 10104159001) and plus lysozyme (Sigma, L6876) plus 1 tablet of protease inhibitor (Sigma, 11836170001). The resuspended pellet was sonicated 2-3x for 5 min (15 sec sonication at 40% output, stop 45 sec, Fischer Scientific model FB120, 0.63cm probe). The lysate was then cleared by centrifuging for 1.5 h at 20,000 x *g* at 4°C. In the meantime, 6 ml of a Ni-NTA slurry was washed with 30 mL of lysis buffer 2x. The clarified lysate was then added to the prewashed Ni-NTA beads and incubated for 1 h rotating at 4°C. The beads were then loaded on polypropylene columns (Qiagen, 34964, 2 columns/L of starting culture), washed two times with 25 ml of lysis buffer and eluted in 4 ml of elution buffer (50 mM Tris-HCl pH 7.5, 150 mM NaCl, 500 mM Imidazole). The eluate was concentrated to ~2.5 ml on a Amicon column (EMD Millipore, UFC901024). The eluate was then buffer exchanged using a PD10 column (according to manufacturer's instructions) into 50 mM Tris-HCl pH 7.5 + 150mM NaCl containing 10% of glycerol, aliquoted, flash frozen and stored at -80°C. Protein purity was assessed by Coomassie staining (Fisher Scientific, SimplyBlue™ SafeStain, LC6060).

**Protein ELISA**

High-binding plates (Corning, 9018) were coated with 0.25 µg of TTFC-SpyTag003 or Diphtheria toxin (Sigma, D0564)/well in PBS overnight at 4°C. The following day, the plates were washed 5x with PBST (PBS + 0.05% Tween-20, Sigma, P1379), blocked with 200 µl of 2% BSA (Sigma-Aldrich, A7030-100G) in PBST at room temperature for 2h. The plates were then incubated with the indicated sera from mice in serial dilutions (typically 1:100 starting dilution, followed by seven 1:3 or 1:4 serial dilutions). Next, the plates were washed 5x with PBST and incubated with secondary antibody at a dilution of 1:10,000 (SouthernBiotech, 1030-05) in 2% BSA/PBST. After five more PBST washes, the plates were developed using 100 µl of TMB reagents (BD, 555214) and the reaction stopped with 50 µl of stop solution (BioLegend, 423001). Antibody titers were calculated by logarithmic interpolation of the dilutions with absorbance at 450 nm immediately above and below the absolute value 0.2, as previously described[4].

**Tetanus toxin challenge experiment**

Tetanus toxin challenge experiments were performed following previously published protocols[8]. Briefly, tetanus toxin (List Biological Laboratories, 190A) was resuspended in 0.9% NaCl plus 1 mg/ml mouse serum albumin (Sigma, A3139), pH 7 (=dilution buffer) at a concentration of 50 µg/ml, aliquoted and stored at -80°C. Because the potency of different toxin preparations can differ widely, we tested a range of toxin concentrations to identify the dose at which mice consistently developed tetani paralysis (i.e., 100 ng/kg). On the experimental day, a frozen aliquot of toxin was thawed to room temperature for 1.5 h and diluted in dilution buffer to 60 pg/µl (150 ng/kg dose) or 44 pg/µl (110 ng/kg dose). For the serum neutralization experiments, the diluted toxin was mixed 1:1 with dilution buffer containing 1:10 of serum from mice colonized with the indicated strain (final concentration of toxin: 22 pg/µl for 110 ng/kg dose) and incubated with rotation at RT for 1h. For the normal challenge experiments, the toxin was mixed 1:1 with dilution buffer and also incubated for 1 h at RT before injection (final concentration of toxin: 30 pg/μl for 150 ng/kg dose). 8-12-week-old C57BL/6 mice were then weighed and injected with either 150 ng/kg (colonized mice experiment) or 110 ng/kg (serum neutralization experiment) of tetanus toxin intraperitoneally (100 μl for a 20 g mouse) as indicated in the figure legends. The mice were monitored for 200 h, the experimental endpoint. Mice were euthanized when they displayed moderate tetanus symptoms (hunched back and disappearance of the righting reflex) and considered positive for tetanus.

**Production of mi3 nanoparticles**

SpyCatcher003-mi3 particles were prepared as previously described[9] by purification from *E. coli* BL21 (DE3) – RIPL competent cells (Agilent) transformed with a pET28a SpyCatcher-mi3 gene comprising an *N*-terminal hexa-histidine tag (Addgene, Plasmid #112255). Briefly, transformed cells were selected for with kanamycin and grown in LB until an $OD_{600}$ of ~0.6-0.8. Protein expression was induced by adding 420 μM IPTG and transferring the cultures to 20˚C, shaking at 200 rpm for 16 h. Cells were harvested by centrifugation (4,000 x *g*), pellets were resuspended in lysis buffer (25 mM Tris-HCl pH 8.0, 300 mM NaCl) plus DNAse (Sigma, 10104159001), lysozyme (Sigma, L6876), and protease inhibitors (Sigma, 11836170001), and lysed with a cell disruptor. Clarified lysates were obtained by centrifugation at 21,000 x *g* for 30 min, and SpyCatcher003-mi3 particles were isolated by ammonium sulfate precipitation (25% saturation), followed by dialysis and size exclusion chromatography (SEC) purification using a HiLoad 16/600 Superose 6 column (Cytiva) equilibrated with 25 mM Tris-HCl pH 8.0 and 150 mM NaCl (TBS). SpyCatcher003-mi3 particles were stored at 4˚C and used for conjugations after filtering with a 0.2-μm filter and centrifugation at 20,000 x *g* for 30 min.

**Preparation of TTFC-mi3 nanoparticles for immunization**

Purified unconjugated SpyCatcher003-mi3 particles were incubated with 3-molar excess of TTFC-SpyTag003 protein at 22ºC for 1 h in TBS. Conjugated particles were separated from unconjugated proteins by SEC on a Superose 6 10/300 column (Cytiva) equilibrated with PBS. Fractions corresponding to conjugated SpyCatcher003-mi3 particles were collected and analyzed by SDS-PAGE. Concentrations of conjugated particles were determined using the absorbance at 280 nm as measured by a Nanodrop spectrophotometer (ThermoFisher). Conjugated particles were stored at 4ºC for up to 1 month after conjugation in PBS buffer. On days 0 and 14, the mice were bled via cheek bleed. For immunization (day 0 and 28), conjugated particles were mixed with 50% v/v PBS/AddaVaxTM adjuvant (Invivogen) (2.5 μg/mouse in 50 μl) and injected intramuscularly in the thigh of preshaved mice. At the experimental endpoint (day 42), the blood was harvested via cardiac puncture and the serum isolated using coagulating tubes (BD, 365967).

### Phylogenetic tree generation

The phylogenetic analysis of *S. epidermidis* strains was generated using kSNP4.1 (https://academic.oup.com/mbe/article/40/11/msad235/7390727), which analyzes single nucleotide polymorphisms across genomes[10]. We used NCBI datasets tool to download the *C. accolens* ATCC 49725 genome from NCBI (accession: GCF_000159115.1, assembly:ASM15911v1). We selected a kmer value of 21 and generated a maximum likelihood tree (full command: kSNP4 -in genome_w_plasmid.list -k 21 -ML -outdir genome_w_plasmid_kSNP_run). The output tree was visualized using iTOL (https://itol.embl.de/). and re-rooted on the *C. accolens* genome to generate the final tree. All the code associated with this analysis can be found here (https://github.com/FischbachLab/pub_djenet_sepi/tree/main/WholeGenomeTree).

### Protein structure predictions using AlphaFold2

To predict the structure of Aap, we submitted the sequence of LM088 Aap to AlphaFold2[11] using (https://neurosnap.ai/). Predicted 3D structures were visualized and rendered using PyMOL v.2.5.5 (Schrödinger, Inc.).

### Statistical analyses

All statistical analyses were performed using the Prism software (GraphPad Software, La Jolla, CA). Paired and unpaired (where appropriate) two-tailed Student's t tests were used when two groups were compared. When more than two groups were compared, a one-way ANOVA analysis was performed with appropriate post-hoc tests: Tukey's when comparing all group combinations or Dunnett's post-hoc test when comparing one group versus all other groups. When data was analyzed over time, a two-way ANOVA was used. When data was plotted on a log10 scale, the statistics were calculated using a log10 transformation of the data. For survival experiments (tetanus challenge), a log-rank test was used. All tests used are indicated in the figure legends and the P values are reported on the figures. P values below 0.05 were considered statistically significant.

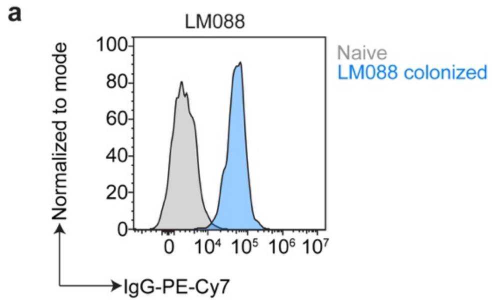


**Figure s1: Colonization with *S. epidermidis* induces a systemic B cell response.** (**a**) *S. epidermidis* strain LM088 was stained with the serum of mice colonized (blue) or not (naïve, grey) with the same strain for 6 weeks and analyzed by flow cytometry. Representative of three independent experiments. Bacteria were gated on Syto9 positive cells.

**a**

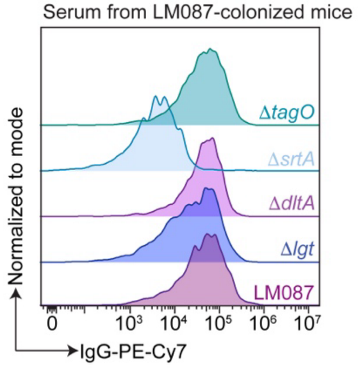


**b**

| LM087 | kDa | LM088 | kDa | |
|---|---|---|---|---|
| FbE | 115 | FbE | 115 | Common substrates |
| OLHLPF_08425 | 81 | DOIKNN_08360 | 85 | |
| OLHLPF_01190 | 75 | DOIKNN_01120 | 75 | |
| OLHLPF_04300 | 41 | DOIKNN_04225 | 40 | |
| OLHLPF_01575 | 21 | DOIKNN_01490 | 22 | |
| | | DOIKNN_00225 | 99 | Single-strain substrates |
| OLHLPF_12100 | 203 | | | |
| | | Aap | 157 | |
| | | DOIKNN_12140 | 22 | |

**Sortase substrates in *S. epidermidis***

**c**

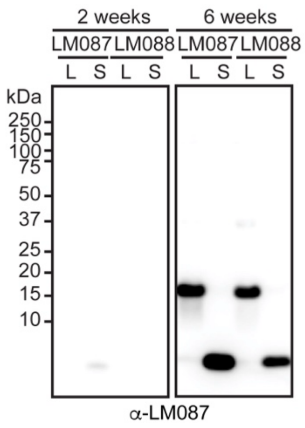


**d**

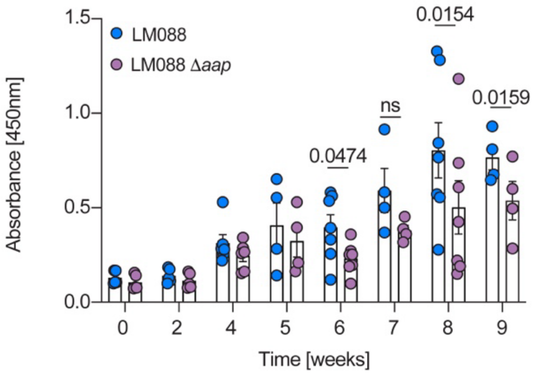


**Figure s2: The B cell response against *S. epidermidis* recognizes the sortase substrate Aap.** (**a**) Flow cytometry analysis using the serum of mice colonized with *S. epidermidis* strain LM087 for 5 weeks against a panel of cell wall mutants of the same strain. One representative of three independent experiments. (**b**) Table of sortase substrates found in LM087 and LM088. (**c**) Immunoblot analysis of LM087 lysate (L) and culture supernatant (S) using the serum of mice colonized with LM087 for 2 or 6 weeks. For the 6 week timepoint, the mice were only colonized every other day for 1 week and not recolonized once/week. One representative of three independent experiments. (**d**) IgA ELISA of the nasal washes of mice colonized with LM088 for the indicated times against LM088 and an Aap-deficient strain of LM088 (Δ*aap*). Of note, the nasal washes were diluted 1:3 and the data for LM088 are the same as the ones displayed on Fig 1. (n=5-9/time point, two independent experiments pooled). The data were analyzed using a paired Student t test at each time point and the graph shows mean +/- SEM.

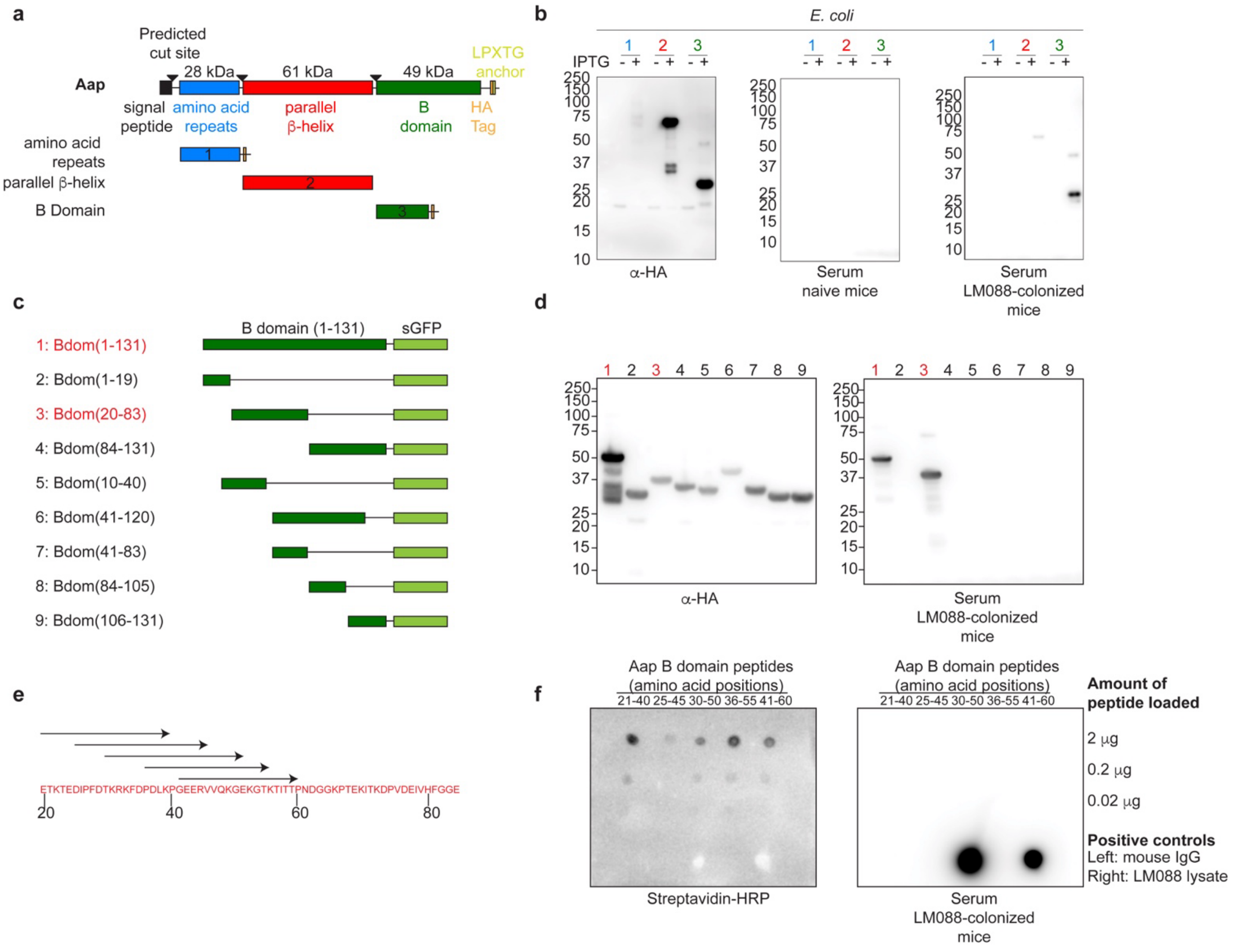


**Figure s3: Identification of the minimal epitope in the B domain of Aap**. (**a**) Schematic of LM088 Aap protein truncations for expression in *E. coli*. (**b**) Immunoblot analysis of the constructs shown in (a) using anti-HA antibody to assess expression (induced by IPTG), using the serum of naïve mice or the serum of mice colonized with LM088. (1= amino acid repeat domain, 2= parallel β-helix domain, and 3=one repeat of the B domain). (**c**) Schematic of the B domain truncations fused to Superfolder GFP (sGFP) for expression in *E. coli*. (**d**) Immunoblot analysis of the truncations shown in (c) using anti-HA antibody or the serum of mice colonized with LM088. (**e**) Minimal epitope identified in the B domain and the peptides synthesized and tested in (f). (**f**) Dot blot analysis of the biotinylated peptides shown in (e) using streptavidin-HRP or the serum of mice colonized with LM088. For all immunoblot analyses, protein ladder sizes are given in kDa. All blots are representative of three independent experiments.

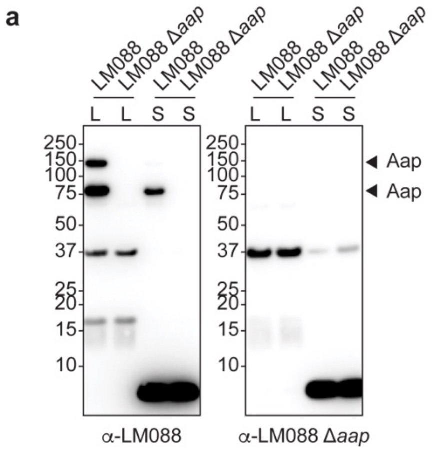


**Figure s4: Immunoblot analysis of mice colonized with LM088 and LM088 Δ*aap***. (**a**) Immunoblot of cell lysate (L) and culture supernatant (S) of LM088 and LM088 Δ*aap* using the serum of mice colonized with LM088 or LM088 Δ*aap* (from the mice shown in Fig. 2j). Representative of two independent experiments.

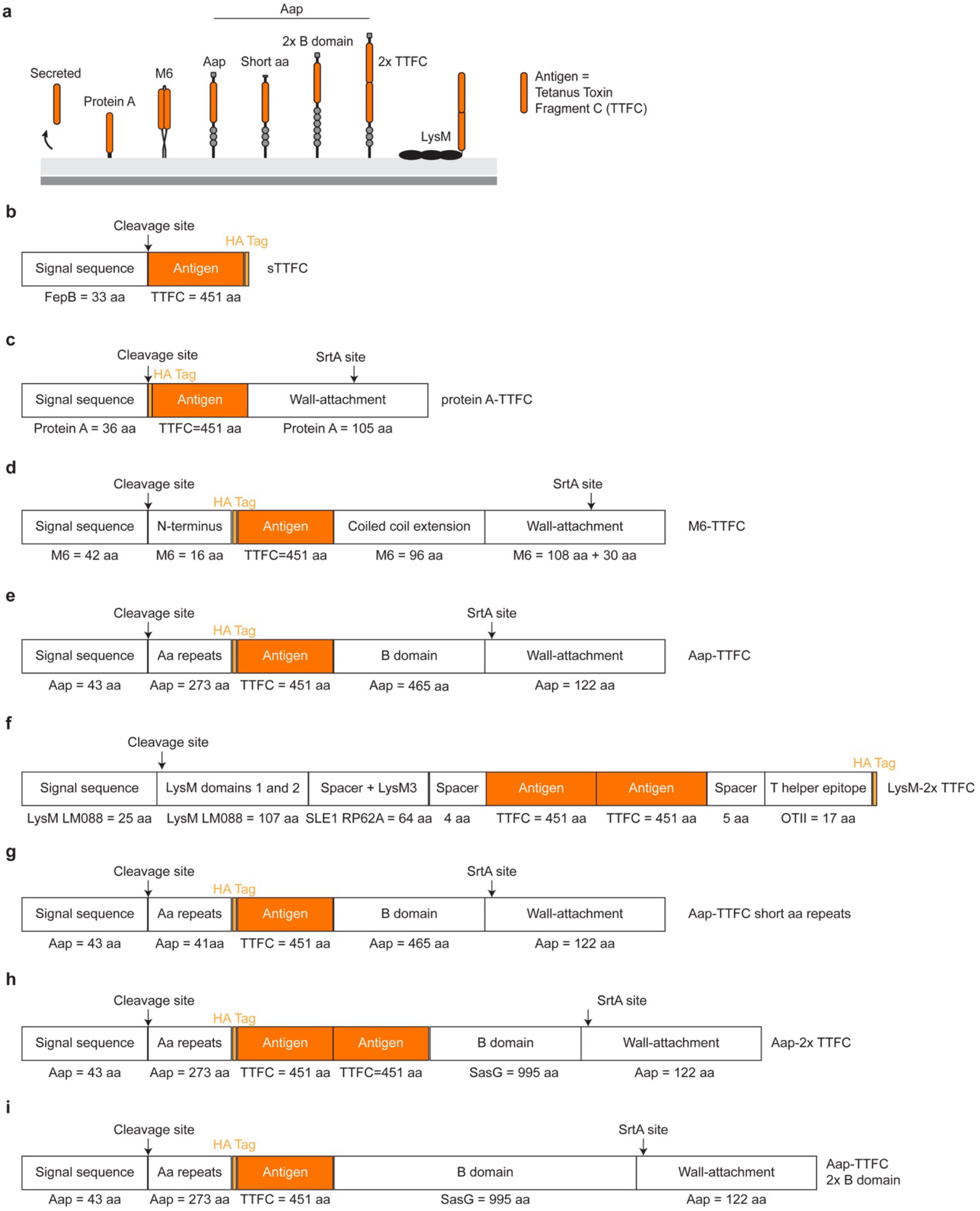

a
Aap
Secreted
Protein A
M6
Aap
Short aa
2x B domain
2x TTFC
LysM
Antigen = Tetanus Toxin Fragment C (TTFC)
b
Cleavage site
HA Tag
Signal sequence
Antigen
sTTFC
FepB = 33 aa
TTFC = 451 aa
c
Cleavage site
HA Tag
SrtA site
Signal sequence
Antigen
Wall-attachment
protein A-TTFC
Protein A = 36 aa
TTFC=451 aa
Protein A = 105 aa
d
Cleavage site
HA Tag
SrtA site
Signal sequence
N-terminus
Antigen
Coiled coil extension
Wall-attachment
M6-TTFC
M6 = 42 aa
M6 = 16 aa
TTFC=451 aa
M6 = 96 aa
M6 = 108 aa + 30 aa
e
Cleavage site
HA Tag
SrtA site
Signal sequence
Aa repeats
Antigen
B domain
Wall-attachment
Aap-TTFC
Aap = 43 aa
Aap = 273 aa
TTFC = 451 aa
Aap = 465 aa
Aap = 122 aa
f
Cleavage site
HA Tag
Signal sequence
LysM domains 1 and 2
Spacer + LysM3
Spacer
Antigen
Antigen
Spacer
T helper epitope
LysM-2x TTFC
LysM LM088 = 25 aa
LysM LM088 = 107 aa
SLE1 RP62A = 64 aa
4 aa
TTFC = 451 aa
TTFC = 451 aa
5 aa
OTII = 17 aa
g
Cleavage site
HA Tag
SrtA site
Signal sequence
Aa repeats
Antigen
B domain
Wall-attachment
Aap-TTFC short aa repeats
Aap = 43 aa
Aap = 41aa
TTFC = 451 aa
Aap = 465 aa
Aap = 122 aa
h
Cleavage site
HA Tag
SrtA site
Signal sequence
Aa repeats
Antigen
Antigen
B domain
Wall-attachment
Aap-2x TTFC
Aap = 43 aa
Aap = 273 aa
TTFC = 451 aa
TTFC=451 aa
SasG = 995 aa
Aap = 122 aa
i
Cleavage site
HA Tag
SrtA site
Signal sequence
Aa repeats
Antigen
B domain
Wall-attachment
Aap-TTFC 2x B domain
Aap = 43 aa
Aap = 273 aa
TTFC = 451 aa
SasG = 995 aa
Aap = 122 aa

**Figure s5: Schematic of the construct designs to redirect the B cell response to *S. epidermidis*.** (**a**) Schematic of the different constructs generated to redirect the B cell response to *S. epidermidis*: (**b**) secreted tetanus toxin fragment C (sTTFC), (**c**) Protein A-TTFC, (**d**) M6-TTFC, (**e**) Aap-TTFC (wTTFC), (**f**) LysM-TTFC, (**g**) Aap-TTFC short amino acid repeats, (**h**) Aap-2x TTFC and (**i**) Aap-TTFC 2x B domain. For more information, including sequences, please see Supplementary Table S1.

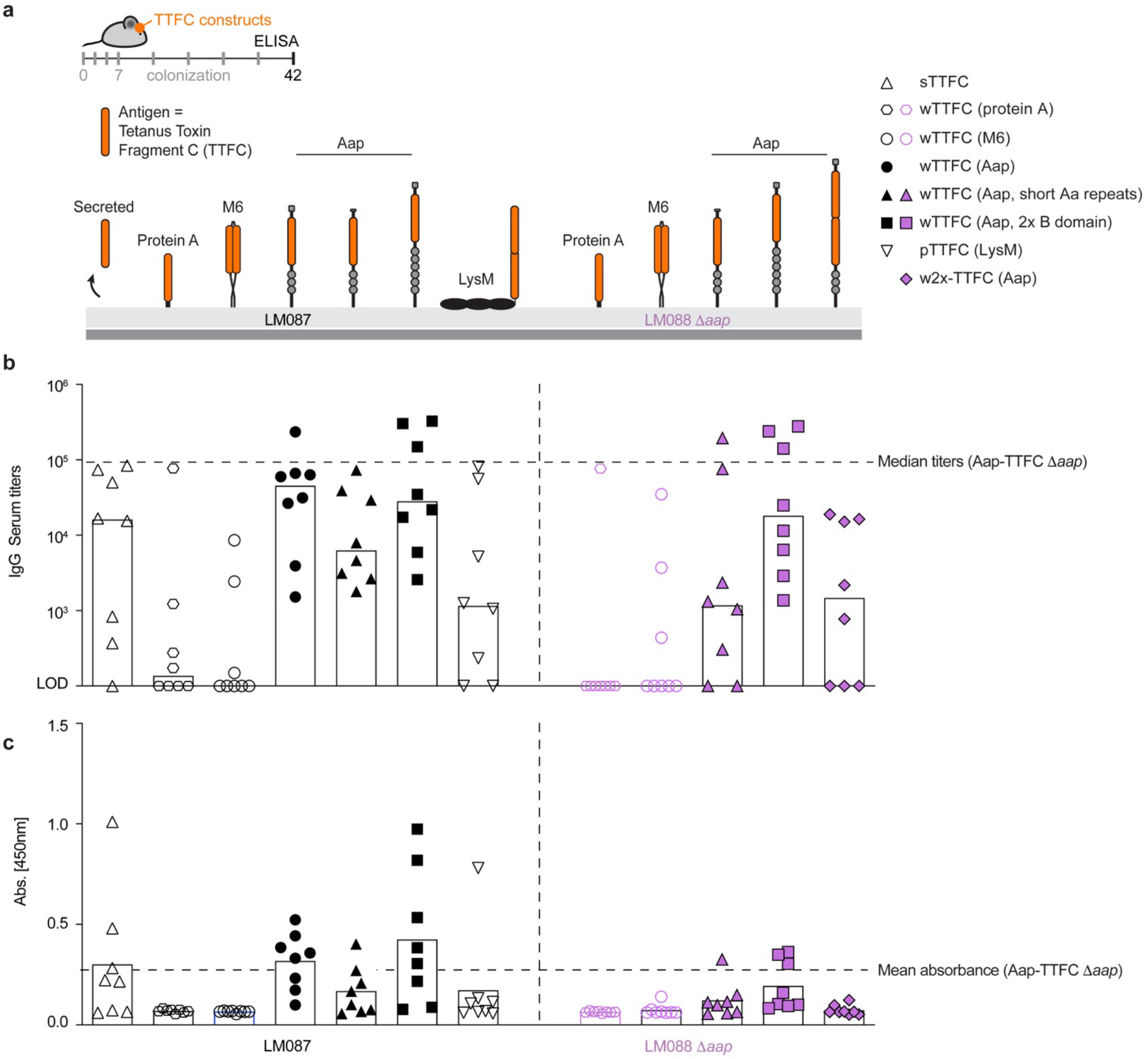


**Figure s6: Redirecting the B cell response to *S. epidermidis* using engineered strains.** (**a**) 6–10-week-old SPF mice were colonized for 6 weeks with *S. epidermidis* strain recombinantly expressing different tetanus toxin fragment C (TTFC) fusion proteins. The constructs were built in either WT LM087 (black) or Δ*aap* LM088 (purple) background strain. (**b**) Serum titers against TTFC 6 weeks post colonization and (**c**) TTFC specific IgA in nasal washes (undiluted washes). N=8/group, two independent experiments pooled. Of note, for clarity panel (b) shows medians and (c) means. The dashed lines show the corresponding median (b) and mean (c) values for strain Δ*aap* + Aap-TTFC (wTTFC) (see Fig. 3d, which was run together with the data shown in fig. s6). p=peptidoglycan targeting (i.e. LysM-2x TTFC). LOD= limit of detection.

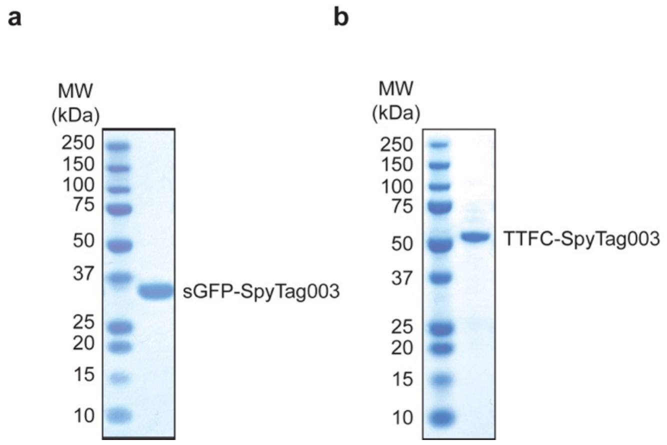


**Figure s7: Purity of proteins purified from *E. coli*.** Coomassie staining of purified superfolder GFP (sGFP)-SpyTag003 (**a**) and tetanus toxin fragment C (TTFC)-SpyTag003 (**b**) proteins. One representative gel of at least three independent experiments.

**Video s1: Mice colonized with wDT are not protected against a lethal dose of tetanus toxin.** Mice were colonized with *S. epidermidis* Δ*aap* expressing Aap-diphtheria toxin (wDT, catalytic mutant of diphtheria toxin) for 6 weeks and challenged with a lethal dose of tetanus toxin (150 ng/kg) (mice shown in **Fig. 3f**). The video was recorded 44 h post injection of the toxin and is representative of two independent experiments.

**Video s2: Mice colonized with wTTFC are protected against a lethal dose of tetanus toxin.** Mice were colonized with *S. epidermidis* Δ*aap* expressing Aap-tetanus toxin fragment C toxin (wTTFC) for 6 weeks and challenged with a lethal dose of tetanus toxin (150 ng/kg) (mice shown in **Fig. 3f**). The video was recorded 44 h post-injection (same experiment/time as mice shown in **Video s1**) of the toxin and is representative of two independent experiments.

**Video s3: Mice colonized with wTTFC do not develop pathology after injection of a lethal dose of tetanus toxin.** Mice were colonized with *S. epidermidis* Δ*aap* expressing Aap-tetanus toxin fragment C toxin (wTTFC) for 6 weeks and challenged with a lethal dose of tetanus toxin (150 ng/kg) (mice shown in **Fig. 3f**). The video was recorded >200 h post-injection (same mice as shown in **Video s2**) and is representative of two independent experiments.
